# Allele-specific gene regulation, phenotypes, and therapeutic vulnerabilities in estrogen receptor alpha mutant endometrial cancer

**DOI:** 10.1101/2022.06.13.495977

**Authors:** Zannel Blanchard, Craig M. Rush, Spencer Arnesen, Jeffery M. Vahrenkamp, Adriana C. Rodriguez, Elke A. Jarboe, Callie Brown, Matthew E. K. Chang, Mark R. Flory, Hisham Mohammed, Katarzyna Modzelewska, David H. Lum, Jason Gertz

## Abstract

Activating estrogen receptor alpha (ER) mutations are present in primary endometrial and metastatic breast cancers, promoting estrogen-independent activation of the receptor. Functional characterizations in breast cancer have established unique molecular and phenotypic consequences of the receptor, yet the impact of ER mutations in endometrial cancer has not been fully explored. In this study, we used CRISPR-Cas9 to model the clinically prevalent ER-Y537S mutation and compared results to ER-D538G to discover allele-specific differences between ER mutations in endometrial cancer. We found that constitutive activity of mutant ER resulted in changes in the expression of thousands of genes, stemming from combined alterations to ER binding and chromatin accessibility. The unique gene expression programs resulted in ER mutant cells developing increased cancer associated phenotypes, including migration, invasion, anchorage independent growth, and growth *in vivo*. To uncover potential treatment strategies, we identified ER associated proteins via Rapid Immunoprecipitation and Mass Spectrometry of Endogenous Proteins (RIME) and interrogated two candidates, CDK9 and NCOA3. Inhibition of these regulatory proteins resulted in decreased growth and migration, representing potential novel treatment strategies for ER mutant endometrial cancer.

**Implications:** This study provides insight into mutant ER activity in endometrial cancer and identifies potential therapies for women with ER mutant endometrial cancer.

**STATEMENT OF SIGNIFICANCE:** Activating estrogen receptor alpha (ER) mutations promote ligand-independent activity of the receptor. This study evaluates ER-Y537S and ER-D538G mutations in primary endometrial cancer, revealing their effects on gene regulation and cancer-associated phenotypes. By identifying ER associated proteins, we also uncover potential novel treatments for women with ER mutant endometrial cancer.

## INTRODUCTION

Estrogen receptor alpha (ER) is a key oncogene in endometrial and breast cancer that acts as a transcription factor when bound to estrogens. The main risk factors for endometrial cancer all implicate excess estrogens, in comparison to progesterone, in promoting the formation of the disease [1]. In breast cancer, the efficacy of hormone therapies such as aromatase inhibitors (AIs), selective estrogen receptor modulators (SERMs), and selective estrogen receptor degraders (SERDs) in ER expressing tumors speak to ER’s fundamental role in disease progression. Consistent with its oncogenic role in both cancer types, activating mutations in the ligand binding domain (LBD) of ER are found in primary endometrial tumors [2–4] and hormone therapy resistant breast cancers [5–9]. These mutations cluster near helix 12 of the LBD, in a region that determines the protein’s conformation and co-activator binding, with most mutations occurring at amino acids Y537 and D538 [5, 7–11].

Studies have indicated that ER mutations are rarely found in primary breast tumors, but are found in approximately 30% of ER-positive metastatic lesions, driving ligand-independent ER activity [5-8, 10, 12-16]. Across multiple *in vitro* models, ER mutations modulate the expression of estrogen responsive genes in addition to unique genes not normally regulated by estrogen [5, 7, 8, 10, 13–16]. ER mutations in breast cancer increase proliferation, migration, and invasion *in vitro* as well as growth *in vivo* [5-7, 10, 12-14]. Multiple preclinical models suggest that ER mutants have decreased affinity for SERMs and SERDs [5, 7, 8, 10, 11, 15]; however, newer SERMs and SERDs, such as lasofoxifene, bazedoxifene, and giredestrant, in addition to strategies that exploit ERs co-regulatory proteins are being explored as potential treatments for patients with ER mutant breast cancer [6, 11, 12, 14, 17–19]. While studies have extensively explored ER mutations in breast cancer models, fewer studies have elucidated their effects in endometrial cancer. The distinction between ER mutations in breast and endometrial cancer is critical, as the molecular responses to estrogen stimulation differ significantly between cancer types [1, 20], potentially through the use of distinct co-regulatory proteins.

In contrast to breast cancer, ER mutations are relatively common in primary endometrial tumors, affecting between 3.7% - 5.5% of endometrial cancer patients each year [2–4, 21]. In endometrial cancer, ER mutations have been identified in patients with lower body mass indexes, indicating a loss of estrogen dependence [3]. To unravel the effects of ER mutations in endometrial cancer, our previous study interrogated the molecular consequences of the ER- D538G mutation in endometrial cancer cells [22]. Cells harboring ER-D538G exhibited ligand-independent ER activity and novel gene regulation mediated by an altered ER binding profile and differential chromatin accessibility [22]. However, it is unclear whether common ER mutations have similar regulatory effects and if the mutations promote more aggressive phenotypes in endometrial cancer. Furthermore, the question of which therapies should be used to treat endometrial cancer patients harboring ER mutant tumors remains unanswered.

We addressed these open questions by modelling the ER-Y537S mutation in endometrial cancer cells and compared the molecular and phenotypic consequences of the ER-Y537S and ER-D538G mutations. We discovered allele-specific changes in the expression of thousands of genes regulated by mutant ER. ER-Y537S mediated gene expression shared similarities with constitutive estrogen signaling, while ER-D538G mediated gene expression was less similar to constant ER stimulation. The changes in gene expression were associated with alterations to ER binding and chromatin accessibility at thousands of genomic loci. ER mutant-specific molecular changes promoted more aggressive cancer associated phenotypes, including an increase in endometrial cancer cell migration, invasion, and anchorage independent growth. ER-Y537S drove increased tumor growth *in vivo* and promoted higher metastatic burden in the lungs after tail vein injection, compared to wildtype endometrial cancer cells. Utilizing Rapid Immunoprecipitation and Mass Spectrometry of Endogenous Proteins (RIME), we identified ER associated proteins in endometrial cancer and explored two candidates as therapeutic targets in ER mutant disease. The studies provide valuable insights into the molecular, phenotypic, and therapeutic vulnerabilities of ER mutant endometrial cancer.

## MATERIALS AND METHODS

### Generation of ER-Y537S cell lines

ER-Y537S mutant cell lines were generated using the CRISPR epitope tagging ChIP-seq (CETCH-seq) method created by Savic et al [23] and previously described in Blanchard et al [22]. In brief, we constructed an ER-Y537S gBlock (Supplemental Table 4) for homology directed repair. Using previously described methods, the ER-Y537S gBlock was cloned into a pFETCH plasmid and transfect along with a plasmid expressing Cas9 and guide RNAs targeting *ESR1* [22]. Ishikawa cells (Sigma-Aldrich) were cultured in full serum media [RPMI 1640 media supplemented with 10% Fetal Bovine Serum (FBS) (Sigma Aldrich), 1% penicillin-streptomycin (Thermo Fisher Scientific)], transfected with aforementioned plasmids and treated with 1 µM SCR7 (Xcessbio) for 72 hours post transfection, to inhibit nonhomologous end joining. Following a media change, transfected cells were treated with 200 µg/mL G418 until resistant cells were identified. Single cells were cultured at a limiting dilution prior to cloning. Colonies were Sanger sequenced and protein expression was validated using FLAG and ER immunoblotting (Supplemental Fig. 1A). Based on expression levels, two ER-Y537S clones were identified for use in future experiments. Two ER-WT and ER-D538G clones established in our prior study were used to establish allele-specific regulatory and phenotypic effects in the current study. Cells were authenticated using STR marker analysis and found to be Mycoplasma negative (IDEXX, tested April 2019).

### Cell culture

Ishikawa cells were purchased from Sigma Aldrich and maintained in RPMI 1640 media (Thermo Fisher Scientific) supplemented with 10% FBS (Sigma Aldrich), 1% penicillin-streptomycin (Thermo Fisher Scientific). For hormone depleted (HD) conditions, cells were cultured in phenol-red free RPMI 1640 media supplemented (Thermo Fisher Scientific) with 10% charcoal-stripped FBS (Sigma Aldrich) and 1% penicillin-streptomycin. For nutrient deprived conditions, cells were cultured in DMEM F12 media (Thermo Fisher Scientific), B-27 supplement (Thermo Fisher Scientific) and Epidermal Growth Factor (EGF) (Thermo Fisher Scientific). All cells were incubated at 37°C in 5% CO_2_.

To visualize ER wildtype and mutant cells in vivo, WT, ER-Y537S and ER-D538G cell lines were transduced with CMV-Luciferase (firefly)-2A-GFP (RFP-Puro) (GenTarget Inc) lentiviral particles. 72 hours post transduction, cells were treated with puromycin at 1 ug/ml until resistant cells remained. Cell lines were maintained in RPMI media with 10% FBS, supplemented with puromycin until they were used in subsequent animal experiments.

### Immunoblotting

Ishikawa parental, ER-WT, ER-D538G, and ER-Y537S cell lines were cultured in 100-mm dishes in full serum media until confluency. Cells were lysed in radioimmunoprecipitation assay (RIPA) buffer (1x PBS, 1% NP-40, 0.5% sodium deoxycholate, 0.1% SDS), supplemented with protease inhibitors (Thermo Fisher Scientific), and briefly sonicated. 40 µg of lysates were loaded on a 4 to 12% Bis-Tris gel (Thermo Fisher Scientific) and immunoblotted with FLAG M2 (Sigma-Aldrich F1804), Estrogen receptor alpha (Millipore Sigma 06-935), CDK9 (Abcam ab239364), and Beta Actin (SantaCruz Biotechnology sc-47778) antibodies were used.

### ERE luciferase reporter assay

ER-WT, ER-Y537S, and ER-D538G cells were seeded in 96-well plates in hormone depleted media for five days. Cells were transfected with dual-luciferase and Renilla ERE reporter vectors (Qiagen), according to the manufacturer’s instructions. 24 hours post-transfection, cells were induced with either 10 nM E2 or DMSO for controls. Luciferase assay was performed with the Dual-Glo luciferase assay kit (Promega) per the manufacturer’s instructions. Experiments were performed in triplicate, with at least two biological replicates. Statistical analysis was performed using a Student’s t-test.

### RNA-seq and analysis

ER-WT, ER-Y537S, and ER-D538G cell lines were cultured in hormone-deprived media for five days and induced with either 10 nM E2 or DMSO for 8 hours. Cells were washed with PBS and lysed using buffer RLT plus (Qiagen) supplemented with 1% beta-mercaptoethanol (Sigma-Aldrich) and passed through a 21-gauge needle and syringe (Sigma-Aldrich). RNA was purified using a Quick-RNA Miniprep kit (Zymo Research) and libraries were generated using the KAPA Stranded mRNA-seq kit (Roche). RNA-seq libraries were sequenced on the Illumina NovaSeq 6000 and sequencing reads were aligned to hg19 build of the human genome using HISAT2 [24]. SAMtools was used to convert SAM files to BAM files [25]. Genes were defined by the University of California Santa Cruz (UCSC) Known Genes and reads that mapped to known genes were assigned with featureCounts [26]. Read counts were normalized and analyzed for differential expression via the DESeq2 package in R [27, 28]. RNA-seq experiments were performed in two ER-Y537S clones treated with DMSO and 10 nM E2, and these clones were evaluated as biological replicates. RNA-seq data from ER-WT and ER-D538G clones generated previously [22] were used as comparisons in differential expression analysis between genotypes. We identified genes that were significantly up- or down-regulated (adjusted P-value <0.05) in response to E2 treatment in ER-WT cells, by comparing ER-WT clones treated with DMSO to ER-WT clones treated with 10 nM E2. In the previous study, we used DESeq2 to compare all ER-D538G samples to ER-WT E2 treated samples and removed ER-WT E2 regulated genes[22]. This resulted in a smaller list of genes regulated by ER-D538G in endometrial cancer cell lines. For this study, we identified ER-Y537S and ER-D538G specific genes by comparing all ER-WT samples to all ER-Y537S and ER-D538G samples, irrespective of DMSO or E2 inductions, which increased power. Genes regulated by E2 in ER-WT cells were then subtracted and the remaining differentially expressed genes with a P-value cutoff of <0.05 were defined as mutant-enriched genes. Mutant-specific gene lists were analyzed using Enrichr Pathway Analysis [29–31] to identify enriched molecular and cellular pathways.

For prolonged E2 RNA-seq experiments, we used gene lists previously defined by treating ER-WT cells with 10 nM E2 every two days with cell lysates collected at day 10, day 15, day 20, and day 25 [22]. RNA-seq libraries were constructed as described above. Differentially expressed genes with an adjusted P-value cutoff of <0.05 were identified by comparing DMSO and 8 hour E2 treated ER-WT cells to 10-, 15-, 20- and 25-day samples.

### Xenograft RNA-seq and analysis

Approximately 30 mg of flash-frozen Ishikawa ER-WT, ER-D538G, and ER-Y537S xenograft tumors taken at the end of the *in vivo* experiments were used for lysis and RNA extraction. Tumor tissue was suspended in 35 uL of lysis buffer per mg of tissue and mechanically homogenized using M tubes (Milteny Biotec) with a gentleMACS Octo Dissociator (Miltenyi Biotec) using program “RNA_02”. Lysis buffer consisted of RLT plus (Qiagen) supplemented with 1% beta-mercaptoethanol (Sigma-Aldrich). After lysis and homogenization, RNA was purified from a volume of lysate equivalent to 15 mg of tissue (525 uL) using a Quick-RNA Miniprep kit (Zymo Research). 500 ng of total RNA was used to generate libraries. Samples were hybridized with Ribo-Zero Gold (Illumina) to deplete cytoplasmic and mitochondrial rRNA. Stranded libraries were prepared using a TruSeq Stranded Total RNA Library Prep Gold kit (Illumina) according to the manufacturer’s instructions. RNA-seq libraries were sequenced on the Illumina NovaSeq 6000. The raw Fastq was filtered for reads specific to human (as compared to mouse) using Xengsort [32]. Fastq files of human-specific reads were processed as described above (RNA-seq and analysis) with mapping to hg19, assigning of reads to UCSC Known Genes, and normalization and analysis of differential expression with DESeq2, as described above. RNA-seq experiments were performed on 3 xenograft tumors for each ER-WT clone 1, ER-D538G clone 1, and ER-Y537S clone 1. Xenografts used were from the *in vivo* xenograft growth study without E2 supplementation. Genes were considered differentially expressed with an adjusted p-value cutoff of <0.05. Differentially expressed gene sets were analyzed using Enrichr Pathway Analysis [29–31].

### ChIP-seq and analysis

ER-WT, ER-D538G, and ER-Y537S cell lines were cultured in hormone-deprived media for five days and induced with either 10nM E2 or DMSO for 1 hour. Cells were fixed with 1% formaldehyde (Sigma-Aldrich) for ten minutes at room temperature. For the CDK9 ChIPs, cells were first fixed with 2 mM disuccinimidyl glutarate (DSG, Thermo Scientific) for 20 minutes at room temperature followed by a second fixation with 1% formaldehyde (Sigma-Aldrich) for 10 minutes at room temperature. 125 nM glycine was added to stop the crosslinking reaction. Cells were washed with cold PBS and chromatin was harvested by scraping the cells in Farnham lysis buffer (5 mM PIPES pH 8.0, 85 mM KCl, 0.5% NP-40) supplemented with protease inhibitors (Thermo Fisher). Chromatin immunoprecipitation was performed as previously described [33] using an anti-FLAG M2 antibody (Sigma-Aldrich F1804) or CDK9 (Abcam ab239364) antibodies. Libraries were sequenced on Illumina NovaSeq 6000 and reads were aligned to hg19 build of the human genome using Bowtie [34] with the following parameters: -m 1 -t --best -q -S -I 32 -e 80 - n 2. SAMtools was used to sort and convert SAM files to BAM files[25]. Peaks with a P-value cutoff of 1 x 10-10 and an mfold parameter between 15 and 100 were called by Model Based Analysis of ChIP-seq-2 (MACS2) [35]. Single fixation input control libraries from ER-WT, ER-Y537S, and ER-D538G were used as controls for FLAG (ER) ChIP-seq experiments and clones were used as biological replicates for each experiment. FLAG (ER) ChIP-seq data from ER-WT and ER-D538G clones generated previously [22] were used as comparisons between genotypes. An input control library was generated by pooling double-fixed chromatin from ER-WT, ER-D538G, and ER-Y537S cells and was used as a control for CDK9 ChIP-seq experiments. CDK9 ChIP-seq was performed on two independent replicates of ER-WT clone 2, ER-D538G clone 1, and ER-Y537S clone 1. DESeq2 [28] was used to normalize read counts and perform differential analysis for FLAG (ER) ChIP at individual ER bound sites. Results from ER-WT cell lines treated with E2 were compared to ER-Y537S and ER-D538G samples treated with DMSO or 10 nM E2. An adjusted P-value cutoff of < 0.05 was used when identifying wildtype and mutant-enriched or depleted ER bound sites. Constant ER binding sites were defined as regions bound in at least one replicate of ER-WT, ER-Y537S, and ER-D538G samples and not differentially bound in the DESeq2 analysis. Similarly, DESeq2 was used to normalize read counts and perform differential analysis for CDK9 ChIP binding. Due to variance in alignment rate among CDK9 ChIP-seq samples, the DESeq2 design formula (∼Alignment_Rate + Genotype) was used where Alignment_Rate was ‘high’ when >50% and ‘low’ when <50% and Genotype reflected ER genotype. An adjusted p-value cutoff of <0.05 was used to identify wildtype and mutant-enriched or depleted CDK9 bound sites. Constant CDK9-bound sites were defined as regions bound by CDK9 in WT E2-treated and mutant DMSO-treated cells but not WT DMSO-treated cells. ERE distribution in constant and mutant-specific sites were calculated by Patser [36], using DNA sequence within 100 bp of peak summits. CDK9 ChIP profile plots were generated using DeepTools [37].

### ATAC-seq and analysis

ER-WT and ER-Y537S cell lines were cultured in hormone-deprived media for five days and induced with either 10nM E2 or DMSO for 1 hour. 250,000 cells per condition were collected for ATAC-seq experiments and subsequent library preparation [38]. Libraries were sequenced on NovaSeq 6000 and aligned to hg19 build of the human genome using bowtie [34] with the following parameters: -m 1 -t --best -q -S -I 32 -e 80 -n 2. SAM files were converted to BAM files and sorted with SAMtools [25]. ATAC-seq peaks were called using MACS2 [35] with an adjusted P-value cutoff of <0.05. FeatureCounts [26] was used to quantify reads that aligned to all ATAC-seq peaks called in any sample. Reads were normalized and analyzed for differential enrichment using the DESeq2 package for R [28]. ATAC-seq experiments were performed in two ER-WT and ER-Y537S clones with a new batch of Tn5 transposase compared to previously published data [22], and clones with the same genotype were used as biological replicates. ER-WT and ER-D538G clones generated previously [22] were used as comparisons in identifying regions of differential chromatin accessibility between genotypes. To identify mutant-enriched and mutant-depleted ATAC-seq sites, all ER-WT samples were compared to ER-Y537S and ER-D538G samples. To identify ER-D538G specific peaks, we compared ER-D538G with ER-WT samples generated with the previous Tn5 transposase batch. To identify ER-Y537S specific peaks, we compared ER-Y537S with ER-WT samples generated with the newer Tn5 transposase batch. To control for differences in the Tn5 transposase batches, we compared older ER-WT to newer ER-WT samples, removing any significantly different peaks from peaks called as significant in the ER-Y537S vs ER-WT and ER-D538G vs ER-WT analyses. BEDTools intersect [39] was used to overlap ATAC-seq regions that were either shared, mutant-enriched, or mutant-depleted with ER bound sites from FLAG ChIP-seq experiments. 500 bp regions surrounding enriched ATAC-seq peaks not associated with ER binding and >2 kb from transcription start sites were utilized for motif analysis using MEME-ChIP [40]. Motifs between 6 and 20 bps in length were identified by MEME ChIP [40], with a motif distribution of zero to one occurrence per sequence.

### Proliferation assays

ER-WT, ER-Y537S, and ER-D538G cells were cultured in full serum or hormone depleted media for three days prior to plating for proliferation experiments. 7500 cells per genotype were seeded in 96-well plates in both media conditions and proliferation was monitored via the IncuCyte Zoom imaging platform (Sartorius) for up to 72 hours, with images acquired every 2 hours. Doubling times for the two clones per genotype in both media conditions were derived from linear regression of the log_2_ confluency percentages normalized to the starting confluency vs. time. Doubling times were calculated as 1 divided by the slope. Experiments were performed in triplicate, with two clones as biological replicates. SI-2 (Cayman Chemicals) and Alvocidib/Flavopiridol (Selleckchem) were used in dose response studies and the calculation of IC_50_ values in ER-WT, ER-Y537S, and ER-D538G cell lines.

### Wound scratch assay

ER-WT, ER-Y537S, and ER-D538G cells were cultured in 96-well ImageLock microplates (Sartorius) in full serum, hormone depleted, and nutrient deprived media until confluent. At confluency, cells were treated with 20 µg/mL Mitomycin C (Sigma-Aldrich) for 4 hours, the monolayer was washed with 1X PBS, scraped, and monitored via live imaging on the IncuCyte Zoom imaging platform (Sartorius) for up to 24 hours, with images acquired every 2 hours. Migration rates were calculated as the relative wound density over time for all cell lines. Experiments were performed in triplicate, with two clones as biological replicates. Statistical analysis was performed using a Student’s t-test. SI-2 (Cayman Chemicals) and Alvocidib/Flavopiridol (Selleckchem) were used in migration studies in ER-WT, ER-Y537S, and ER-D538G cell lines.

### Transwell invasion assay

Cell invasion assays were performed using the 24-well Corning BioCoat Matrigel Invasion Chambers with 8.0 µm pores (Thermo Fisher Scientific). ER-WT, ER-Y537S, and ER-D538G cells were seeded in hormone depleted media for three days prior to plating. At 72 hours, 20,000 cells per genotype were placed in the upper chamber containing phenol red free RPMI1640 (Thermo Fisher Scientific). The bottom chamber contained phenol-red free RPMI 1640 media supplemented with 1% penicillin-streptomycin and 10% FBS as a chemoattractant. Cells were incubated at 37°C with 5% CO_2_ for 48 hours. Invasive cells on the bottom of the chamber inserts were washed with 1X PBS, fixed with 3.7% formaldehyde for 15 minutes and stained with 0.5% crystal violet for 20 minutes at room temperature. Invasive cells were visualized via a stereomicroscope (Nikon SMZ18; magnification, x10) and measured by counting the number of cells in four randomly selected high-power fields located across the center and periphery of the membrane using ImageJ software. All experiments were performed in triplicate, with two biological replicates. Statistical analysis was performed using a Student’s t-test.

### Soft agar colony formation assay

Anchorage-independent growth was evaluated by the soft agar colony formation assay in ER-WT, ER-Y537S, and ER-D538G cell lines. 20,000 cells in single cell suspension per genotype were seeded in 6-well plates containing a 0.5% bottom layer, 0.3% top layer of agar and 1.5 mL of full serum media. Cells were cultured for approximately 27 days with weekly media changes. At the experiment’s endpoint, colonies were stained with 1 mg/mL nitro blue tetrazolium chloride (NBT) overnight and imaged with the Azure biosystems C600 imaging system. Average colony number was measured using ImageJ software, experiments were performed in triplicate with two clones as biological replicates.

### Rapid Immunoprecipitation and Mass Spectrometry of Endogenous proteins (RIME)

RIME experiments were performed as previously described [41]. Briefly, Ishikawa cells were cultured in 15-cm dishes in hormone depleted media for five days prior to a 1 hour 10 nM E2 induction. Cells were fixed in 1% formaldehyde, 0.1 M NaCl, 1 mM EDTA pH 8.0 and 50 mM HEPES ph 7.9 for 15 minutes at room temperature. Fixation was stopped by the addition of 1.1 mL of 2.5 M glycine solution for five minutes at room temperature. For Ishikawa parental cells, the fixed cells were first washed in 10 mL of 0.5% Igepal-PBS, followed by a second wash with 0.5% Igepal-PBS supplemented with 100 µl phenylmethanesulfonyl fluoride (PMSF). Cells were then snap frozen, samples were stored in -80°C until shipment to Active Motif to undergo nuclei isolation, sonication, immunoprecipitation (ER and control IgG), and protein identification via mass spectrometry. Spectra analysis was performed using the Scaffold software and TandemX! Scoring.

For ER mutant Ishikawa cells, crosslinked cells underwent our standard ChIP-seq protocol through immunoprecipitation. Samples were then washed 6 times in RIPA buffer and twice with 100 mM ammonium hydrogen carbonate before bead pellets were frozen at -80°C. Tryptic digest was performed as previously described [41] and peptides obtained were run on the Thermo Scientific Q-Exactive High-Field Orbitrap mass spectrometry instrument coupled to a Thermo Scientific UltiMate 3000 RSLCnano for nanoflow liquid chromatography (LC) in the Oregon Health and Science University (OHSU) Proteomic Shared Resource Core. A standard top 10 data-dependent acquisition for label-free shotgun analysis was employed over the course of a total 90-minute one-dimension LC analysis with the 5-35% portion of the acetonitrile gradient lasting approximately 60 minutes. Raw data was searched using MaxQuant v1.6.17.0 against the UniProt human database from August 29, 2019. Protein and PSM FDRs were set to 0.01. Variable modifications were Oxidation (M), Acetyl (Protein N-term), and Deamidation (NQ). Match between runs (MBR) was enabled. MaxLFQ was enabled with the minimum ratio count set to 1. MS1 precursor intensities were used for relative quantitation. Processed data was further analyzed in Perseus with log2-transformed intensities normalized by median scaling. Missing values were imputed from a normal distribution 0.8 standard deviations wide and centered 1.3 standard deviations away from the protein intensity distribution.

ER pulldown peptide counts in breast cancer cell lines were obtained from previously published datasets [42]. DESeq2 [28] was used to compare peptide counts of ER pulldowns to IgG pulldowns in Ishikawa cell lines. The data was then compared to published peptide counts in MCF-7 breast cancer cell lines [42]. Data normalization and differential enrichment analysis was also performed using DESeq2. Enrichment thresholds of 1.5 log2 fold change over IgG with adjusted p-values of < 0.05 and an 80% confidence in peptide match were applied.

### Immunohistochemistry (IHC)

Tissue samples were fixed in 10% neutral buffered formalin for 24 hours, then washed 3x in 1X PBS. Tissues were then transferred to 70% ethanol and submitted to ARUP Laboratories for histological preparation and H&E staining. In brief, samples were embedded in paraffin and 5 μm sections were prepared. Sections were stained with hematoxylin and counterstained with eosin. H&E slides were analyzed for morphological features and scored by certified pathologist Dr. Jarboe. Ki-67 immunohistochemistry (IHC) was performed on 5 μm sections using a mouse on mouse immunodection kit (Vector Laboratories BMK-2202). Briefly, antigen retrieval was performed using 10 mM sodium citrate pH 6.0 for 20 minutes, endogenous peroxidase blocking was performed by incubation in 3% hydrogen peroxide in methanol, and blocking was performed for 1 hour using M.O.M. blocking reagent (Vector Laboratories BMK-2202), avidin, mouse FcR blocker (Miltenyi Biotec 130-092-575), added to a base blocking buffer consisting of 10% normal goat serum, 5% BSA and 0.5% Tween 20 in 1xPBS. Incubation with primary Ki-67 antibody (Cell Signaling Technology 4494) at 1:400 in M.O.M diluent supplemented with biotin (Vector Laboratories BMK-2202) overnight at 4°C. Incubation with secondary antibody was performed using a 1:250 dilution of biotinylated anti-mouse IgG in M.O.M. diluent (Vector Laboratories BMK-2202) at room temperature for 30 minutes. Avidin/Biotinylated Enzyme Complex (ABC) (Vector Laboratories PK-4000) and DAB (Vector Laboratories SK-4100) development were performed according to manufacturer’s instructions, with ABC incubation for 30 minutes and DAB incubation for 5 minutes. Slides were then counterstained with hematoxylin prior to mounting.

Ki-67 quantification was performed using QuPath (v0.4.3) [43] on digitized slides on one tumor from each genotype matching one of the replicates used in the xenograft RNA-seq experiments. Stain vectors for DAB (Ki-67) were estimated for each slide and Ki-67 positivity was assessed in fifteen 250 μm x 250 μm fields per tumor. Fields were selected to avoid regions of necrosis. The following parameters were used to identify Ki-67 positive nuclei; detection image:optical density sum, requested pixel size:0.4 μm, background radius:8 μm, use opening by reconstruction:checked, background radius:1.5 μm, sigma:1.5 μm, minimum area:10 μm^2^, maximum area:350 μm^2^, threshold:0.02, max background intensity:3, split by shape:checked, exclude DAB (membrane staining):unchecked, cell expansion:5 μm, include cell nucleus:checked, smooth boundaries:checked, make measurements:checked, and score compartment:nucleus DAB OD mean. Cells were considered positive if the nucleus DAB OD mean was > 0.5.

### Animal experiments

All animal experiments were approved by the University of Utah Institutional Animal Care and Use Committee. A total of 60 mice (6-9 weeks old; 17-27 g) were used in the *in vivo* xenograft growth study. A total of 78 mice (5-9 weeks old; 17-27 g) were used in the SI-2 and TP-1287 drug inhibitor study. Mice were housed in a temperature and humidity-controlled room with 12 hour light/dark cycles, and ad libitum access to standard rodent food and water throughout the study duration.

For the growth study, six- to nine-week-old ovariectomized NRG mice were implanted with beeswax pellets containing 0.57 mg E2 under the skin on the back, between the shoulder blades. Half of the cohort received E2 pellet supplementation. Mice were then injected into the right mammary fat pad with either 1x10^6^ ER-WT, ER-Y537S, or ER-D538G cells to create six experimental groups of mice: E2 supplemented wildtype clones (n=10), E2 supplemented Y537S clones (n=10), E2 supplemented D538G clones (n=10), no E2 supplemented wildtype clones (n=10), no E2 supplemented Y537S clones (n=10), and no E2 supplemented D538G clones (n=10). Thirty-four days after cell injection, mice were sacrificed, tumors were excised and weighed. Tumor volume was calculated using the following equation: Volume = (Length x Width^2^)/2.

For the xenograft treatment study, five- to nine-week-old ovariectomized NRG mice were implanted with beeswax pellets containing 0.57 mg E2 in the left subcutaneous space. All mice received E2 pellet supplementation. Mice were then injected into the left flank with either 1 x 10^-6^ ER-WT, ER-Y537S, or ER-D538G cells to create nine experimental groups of mice: E2 supplemented wildtype clones (n=10), E2 supplemented ER-Y537S clones (n=10), E2 supplemented ER-D538G clones (n=10), E2 supplemented wildtype clones treated with 5 mg/kg SI-2 (n=8), E2 supplemented ER-Y537S clones treated with 5 mg/kg SI-2 (n=8), E2 supplemented ER-D538G clones treated with 5 mg/kg SI-2 (n=8), E2 supplemented wildtype clones treated with 10 mg/kg TP-1287 (SMP Oncology) (n=8), E2 supplemented ER-Y537S clones treated with 10 mg/kg TP-1287 (n=8), and E2 supplemented ER-D538G clones treated with 10 mg/kg TP-1287 (n=8). Control mice were treated with PBS via oral gavage once a week or intraperitoneal (IP) injection five days a week. SI-2 treated mice were injected with 5 mg/kg of drug five days a week via IP delivery. TP-1287 treated mice were given with 10 mg/kg of drug via oral gavage once a week. Mice were imaged by IVIS (Perkin Elmer IVIS Spectrum In-Vivo Imaging System) and at the experiment’s endpoint animals were sacrificed, tumors and lungs were excised and weighed. Tumor volume was calculated using the following equation: Volume = (Length x Width^2^) /2.

For the in vivo metastasis study, six- to nine-week-old ovariectomized NRG mice were injected in the tail vein with 1x10^6^ ER-WT, ER-Y537S and ER-D538G cells to create three experimental groups of mice: wildtype clones (n=10), ER-Y537S clones (n=10) and ER-D538G clones (n=10). Mice were injected with 200 μL Luciferin (16.67 mg/ml; GoldBio) 10 minutes before weekly IVIS imaging. Images of the whole mouse tumor burden are presented as log counts of radiance per genotype. Quantification of luminescence is presented as total flux (photons/second) per genotype. Statistical analysis was performed using a Mann-Whitney U tests due to non-normality.

### Statistical analysis and graphical packages

Statistical analyses were performed in R version 3.5.2 (R Core Team 2017), except for P-values for gene pathway analysis, which were calculated by Enrichr [29–31] and P-values for enriched motifs calculated by MEME suite [40]. Heatmaps for gene expression and differential chromatin were generated in R, using the pheatmap package. ChIP-seq heatmaps were generated by displaying the Z-score based on reads per million that aligned to each region using DeepTools [37]. Statistical tests used and P-values can be found throughout the text.

### TCGA data analysis

RNA-seq, mutation data, and clinical data was obtained from the TCGA data portal in December 2015. For survival analysis, the gene expression measurements used were level 3 RNAseqV2 normalized RSEM data. Only samples with endometrioid histology were analyzed. Cox regression was used to statistically analyze the association between gene expression and progression-free survival in R using coxph from the survival package. We used a median cutoff to define high and low expressing tumors for each gene when running the Cox regression. For differential expression analysis between ER wildtype and ER mutant tumors, TCGA samples with the histological type Uterine Endometrioid Carcinoma were selected. Using mutation information taken from cBioPortal [44, 45], those samples with mutations at the amino acids 537 and 538 in the ESR1 gene were compared to those who lacked any mutations in the ESR1 gene. Differential expression analysis was done using DESeq2.

### Data access

All raw and processed sequencing data generated in this study are available under the NCBI Gene Expression Omnibus (GEO; http://www.ncbi.nlm.nih.gov/geo/) accession numbers: GSE205878 (RNA-seq), GSE231744 (xenograft RNA-seq), GSE205877 (ChIP-seq), and GSE205875 (ATAC-seq).

## RESULTS

### ER-Y537S exhibits ligand-independent transcriptional activity in endometrial cancer cells

We generated heterozygous ER-Y537S mutant clones using a strategy we published previously [22] in Ishikawa cells (Supplemental Fig. 1A), a cell line model for ER-positive endometrial cancer with an ER binding profile similar to patient tumors [46]. We initially assessed ER transcriptional activity via a luciferase estrogen response element (ERE, ER’s preferred DNA binding sequence) reporter assay, comparing ER-Y537S clones to previously generated Ishikawa wildtype and ER-D538G cell lines (Fig. 1A). We observed increased ER activity in estrogen depleted conditions for ER-Y537S and ER-D538G cells as compared to wildtype controls. Estrogen induced ER mutant lines showed an increase in luciferase signal, with levels comparable to estradiol (E2) treated wildtype cells. These experiments confirm the ligand-independent transcriptional activity of ER mutants in endometrial cancer cells. Additionally, the creation of ER mutant models allowed us to systematically profile the most clinically prevalent mutations in endometrial cancer.

**Figure 1.**
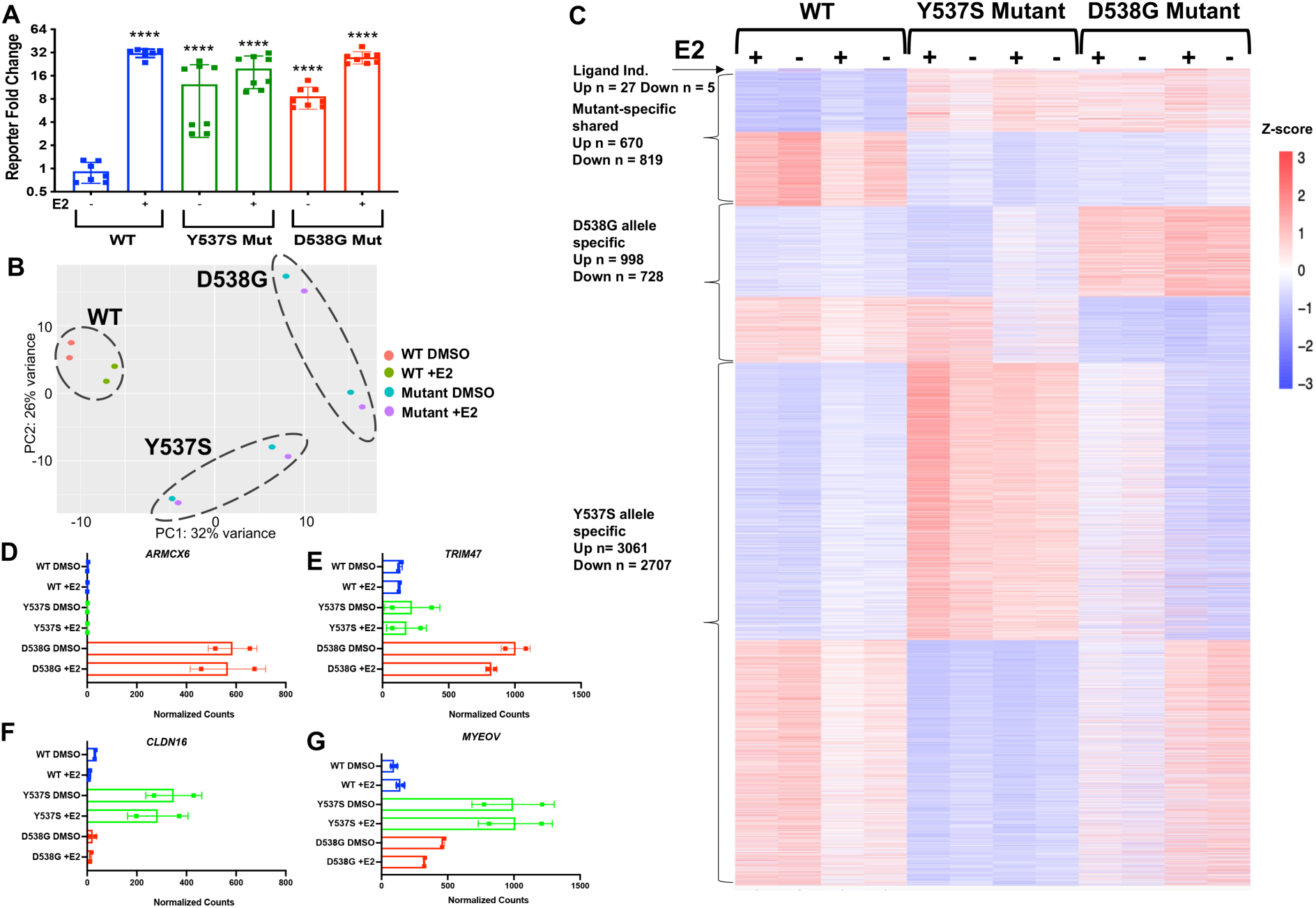
Mutant ER regulates gene expression in a mutation-specific manner in endometrial cancer. (A) ER activity in wildtype, ER-Y537S and ER-D538G cell lines, was measured by ERE driven luciferase activity. ERE luciferase experiments were performed in triplicate, data represents the average luciferase activity for 2 wildtype, 2 ER-Y537S, and 2 ER-D538G cell lines treated with DMSO and E2. Indicated p-values are relative to wildtype without E2. (B) Principal component analysis of RNA-seq data shows the association between wildtype, ER-Y537S, and ER-D538G cell lines. (C) Heatmap displays relative expression levels of ligand independent, mutant-specific shared, ER-D538G, and ER-Y537S specific genes. RNA-seq normalized counts of ER-Y537S regulated genes CLDN16 (D) and MYEOV (E) and ER-D538G regulated genes ARMCX6 (F) and TRIM47 (G). Error bars represent SEM.

### ER-Y537S and ER-D538G cause mutation-specific gene expression changes in endometrial cancer

To understand the allele-specific genome-wide transcriptional changes mediated by ER mutations, we performed RNA-seq on the ER-Y537S clones cultured in hormone depleted media and stimulated with either DMSO or 10nM E2 for 8 hours. We then compared the ER-Y537S data to our previously published wildtype and ER-D538G data [22] to gain a comprehensive understanding of how common ER mutations affect gene expression in endometrial cancer cells. Principal component analysis of the samples clustered wildtype clones together; however, within this cluster, samples segregated by treatment conditions (Fig. 1B). ER-Y537S and ER-D538G clones clustered separately regardless of treatment, with differences between clones becoming apparent. The first principal component (PC1) separated samples based on ER mutation status, clearly separating wildtype, ER-Y537S, and ER-D538G samples and suggests that the mutations are clearly distinct despite their proximity within helix 12 of ER’s ligand binding domain.

When comparing genes regulated by E2 in wildtype cells, 32 genes were up- (27) or down- (5) regulated and shared by both ER mutations (adjusted P-value <0.05; Fig. 1C; for gene lists, see Supplemental Table 1). This demonstrates mutant ER’s ability to regulate the expression of some estrogen-responsive genes and confirms the ligand-independent activity of both mutant receptors [22]. In addition to E2 regulated genes, ER-Y537S and ER-D538G regulated the expression of thousands of genes that were not induced with the transient 8 hour E2 treatment in wildtype cells. We identified 7257 genes differentially up- (3731) and down- (3526) regulated by ER-Y537S in Ishikawa cells (examples in Fig. 1D, E) and 3215 genes differentially up- (1668) and down- (1547) regulated by ER-D538G (examples in Fig. 1F, G). Further comparison of mutation-specific profiles indicated 46% of ER-D538G and 20% of ER-Y537S regulated genes were shared between the mutations, suggesting large ER mutant allele-specific effects, as previously shown in breast cancer models [13, 14].

To determine if these gene expression changes can be observed in patient samples, we analyzed gene expression data from The Cancer Genome Atlas (TCGA) [4]. We identified 230 genes that were differentially expressed between ER wildtype and ER mutant endometrial tumors, using only tumors with endometrioid histology. The 230 genes overlapped significantly with genes regulated by mutant ER in our models. ER-Y537S impacted genes overlapped strongest with differentially expressed genes in patient tumors (4.8-fold enrichment – P-value = 5.1 x 10^-9^, up genes; 1.7-fold enrichment – P-value = 1.2 x 10^-4^, down genes); however, ER-D538G regulated genes were also significantly enriched in the patient differentially expressed genes (3.1-fold enrichment – P-value = 0.0019, up genes; 1.6-fold enrichment – P-value = 0.0015, down genes). These results indicate that ER mutant gene expression patterns found in models can also be observed in endometrial tumors.

To identify potential phenotypes being altered in ER mutant endometrial cancer, we used pathway analysis to identify molecular and cellular functions enriched in genes differentially regulated by ER-Y537S and ER-D538G. ER-Y537S was associated with increased mRNA binding and protein serine/threonine kinase activity (Supplemental Fig. 1B) with ER-D538G associated with increased phospholipase and histone demethylase (H3-K9 specific) activity (Supplemental Fig. 1C). Both mutations showed downregulation of genes involved in actin and cadherin binding, which studies suggest are consistent with more aggressive endometrial disorders [47] and could contribute to tumor progression [48]. The gene ontology findings from our studies in endometrial cancer are consistent with changes observed in ER mutant breast cancer studies. To investigate gene expression differences between cancer types, we compared genes differentially regulated by ER mutations in endometrial cancer cells with genes consistently shared between multiple ER mutant studies done in MCF-7 and T47D cell lines [13–15]. When comparing Ishikawa and T47D cell lines, ER-Y537S regulated transcriptomes showed 35% overlap of up-regulated genes (P-value = 2.3 x 10^-186^, hypergeometric test) and 27% overlap between down-regulated genes (P-value = 9.2 x 10^-88^, hypergeometric test). We observed a similar trend when comparing Ishikawa and MCF-7 ER-Y537S regulated genes, with 38% overlap of up-regulated genes (P-value = 3.2 x 10^-207^, hypergeometric test) and 35% of down-regulated gene sets (P-value = 1.97 x 10^-167^, hypergeometric test). In contrast, ER-D538G regulated genes in Ishikawa and T47D cell lines shared 14% overlap of up-regulated (P-value = 8.6 x 10^-6^, hypergeometric test) and 16% overlap of down-regulated genes (P-value = 2.3 x 10^-11^, hypergeometric test). ER-D538G up-regulated genes in mutant Ishikawa and MCF-7 cell lines showed an 7% overlap (P-value = 0.16, hypergeometric test), with down-regulated genes showing an 8% overlap (P-value = 1.3 x 10^-3^, hypergeometric test). The numbers indicate ER-D538G regulated genes have less overlap between endometrial and breast cancer cell lines. ER-Y537S demonstrated an increased overlap of mutant-regulated genes between cancer types, in addition to being more similar to sustained estrogen signaling (see below). The findings suggest ER mutations regulate different gene expression programs between cancer types, although there are shared, mutation-specific genes potentially regulating similar processes [13, 14].

Our previous publication showed a significant association between ER-D538G regulated genes and worse endometrial cancer patient outcomes, by analyzing disease-free survival in high ER expressing tumors with endometrioid histology in the TCGA endometrial cancer cohort [22]. We queried the connection between patient outcomes and ER-Y537S specific gene expression and found significant overlaps, with ER-Y537S up-regulated genes having a 1.7-fold enrichment for genes associated with poorer outcomes (P-value = 6.2 x 10^-11^, hypergeometric test) and ER-Y537S down-regulated genes showing a 1.42-fold enriched for genes associated with better disease-free survival (P-value = 1.1 x 10^-3^, hypergeometric test) (Supplemental Fig. 1D). In comparison, ER-D538G up-regulated genes showed a 1.5-fold enrichment for genes associated with poorer outcomes (P-value = 8.4 x 10^-3^, hypergeometric test) and down-regulated genes were 1.72-fold enriched for genes associated with better outcomes (P-value = 3.6 x 10^-4^, hypergeometric test) (Supplemental Fig. 1E). This data suggests the identification of an ER mutant gene expression signature that is correlated with worse endometrial cancer patient outcomes, but also emphasizes how findings from cell line models can support and inform future clinical studies.

### Gene regulation by ER-Y537S is similar to constitutive estrogen signaling

The number of genes regulated by ER-Y537S independent of an estrogen induction suggested a significantly altered transcriptional network in Ishikawa cells. These changes could be the result of novel functions attributed to the mutant receptor or to its constitutive activity. To delineate these changes, we compared ER-Y537S regulated genes to our previously published data from wildtype Ishikawa cells treated with 10nM E2 for up to 25 days [22]. The majority (62%, n = 406) of genes up-regulated in response to prolonged E2 treatment in wildtype cells overlapped with ER-Y537S specific up-regulated genes (P = 1.98 x 10^-180^; hypergeometric test) (Supplemental Fig. 2A; for gene lists, see Supplemental Table 2) while 71% (n = 806) of genes down-regulated in response to prolonged E2 treatment in wildtype cells overlapped with ER-Y537S specific down-regulated genes (P = 1.67 x 10^-462^; hypergeometric test) (Supplemental Fig. 2B). This pattern of ER-Y537S mediated gene expression is distinct from ER-D538G regulation, where only 11% of prolonged E2 up- and down-regulated genes overlapped with the ER-D538G regulated genes (up - 72, P = 4.2 x 10^-6^; hypergeometric test; down – 122, P = 6.79 x 10^-11^; hypergeometric test) (Supplemental Fig. 2C - D; for gene lists, see Supplemental Table 2). The significant overlaps between prolonged estrogen signaling and ER-Y537S mediated gene regulation suggest that the effects of ER-Y537S are partially explained by constant activity; however, thousands of ER-Y537S regulated genes are not impacted by prolonged estrogen treatment.

### ER-Y537S and ER-D538G mutations significantly expand ER binding

To understand the underlying causes of mutant mediated gene expression, we hypothesized that changes in ER binding could be a contributing factor. To test this hypothesis and elucidate how mutations affect ER genomic binding, we performed Chromatin Immunoprecipitation sequencing (ChIP-seq) of FLAG-tagged ER-Y537S cell lines, cultured in hormone deprived media for 5 days and treated with DMSO or 10nM E2 for 1 hour. The ER-Y537S binding data was then compared to published data from wildtype and ER-D538G mutant cells under identical conditions [22]. ChIP-seq analysis identified 5815 genomic sites bound by ER-Y537S and 4460 sites bound by ER-D538G in Ishikawa cells. In the absence of an estrogen induction, ER-Y537S and ER-D538G were recruited to 3284 sites that were shared with the wildtype receptor after an E2 induction (constant ER binding sites) (Fig. 2A). We found that 2531 sites exhibited increased ER ChIP-seq signal in ER-Y537S samples compared to wildtype, while 2463 sites showed decreased signal. In contrast, 1176 ER-D538G sites showed increased ER ChIP-seq signal compared to wildtype, while 974 regions showed decreased signal (Supplemental Fig. 3A). Genomic location analysis suggested that ER-Y537S specific sites were enriched for promoter regions (P-value < 2.2 x 10^-16^; odds ratio = 1.82, Fisher’s Exact test), while ER-D538G specific sites were enriched for promoter-distal regions (P-value < 2.2 x 10^-16^; odds ratio = 19.8, Fisher’s Exact test). This data suggests that ER-Y537S and ER-D538G bind to the genome independent of ligand and with altered genomic binding site selection. The relative amount of differential genomic binding was different between ER mutations, with ER-Y537S mediating increased ER recruitment, which correlated with larger gene expression changes compared to ER-D538G.

**Figure 2.**
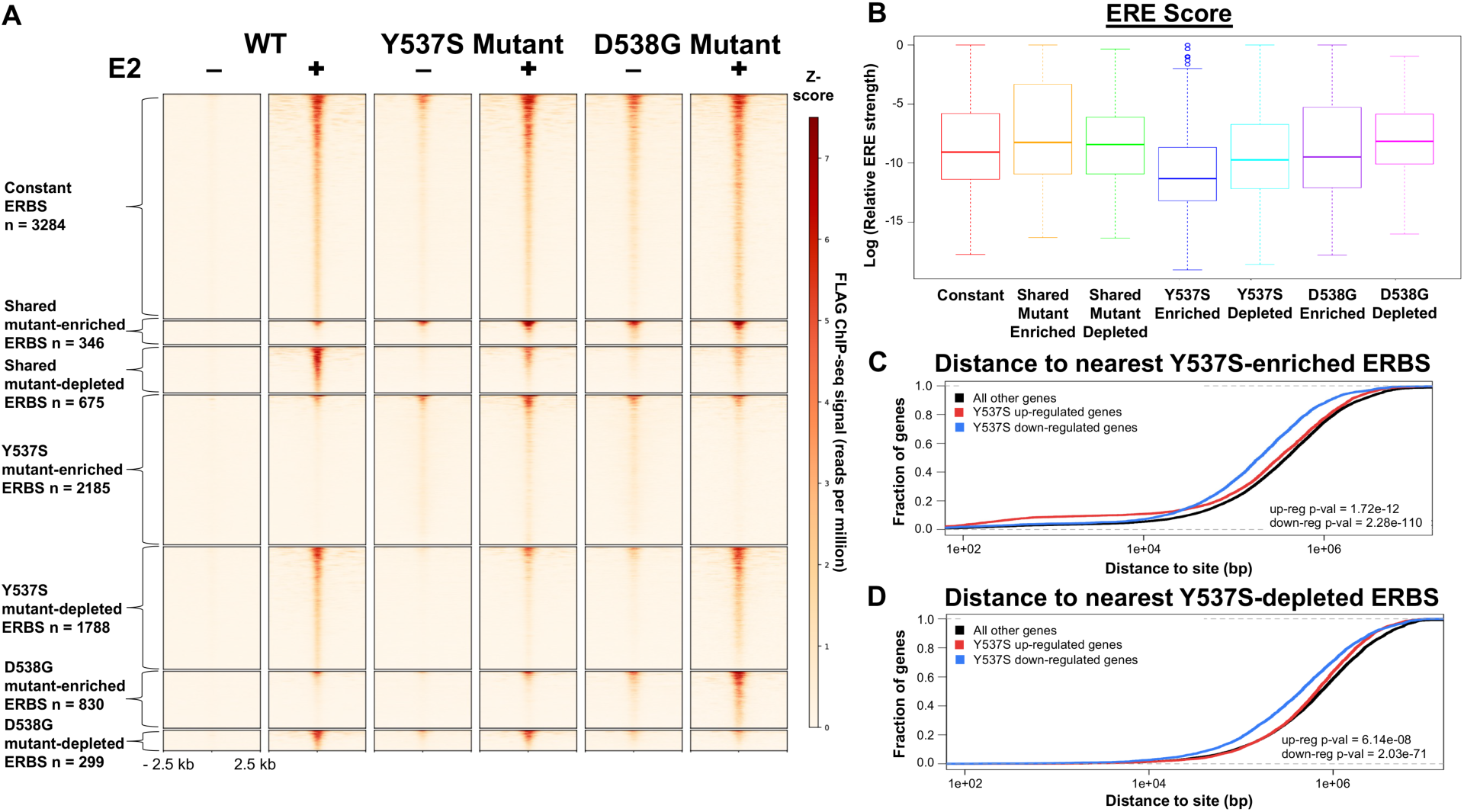
Mutant ER significantly reprograms ER genomic binding. (A) Heatmap shows ER binding patterns in wildtype, ER-Y537S, and ER-D538G cell lines. Figure includes sites that are constant in wildtype and mutant cell lines, sites that are shared between mutations, and sites that are enriched in ER-Y537S and ER-D538G. (B) Relative affinity for ER as measured by the best match to an optimal Estrogen Response Element (ERE), is displayed for each category of ER bound sites. Cumulative distance plots show the fraction of ER-Y537S enriched up-regulated, down-regulated and not regulated genes that have an ER-Y537S enriched (C) or ER-Y537S depleted (D) ER bound site (ERBS) within a specified distance from their transcription start sites.

ERE analysis revealed that the canonical ER binding motif was enriched in constant and shared mutant enriched sites compared to ER-Y537S and ER-D538G enriched ER bound sites (P-value < 2.2 x 10^-16^; Wilcoxon rank sum test) (Fig. 2B). The presence of weaker EREs in ER-Y537S enriched sites is similar to what was observed with ER-D538G enriched binding, confirming that mutant ER is capable of binding regions with less optimal EREs [22]. To further understand how ER binding relates to gene expression changes, we evaluated the distance between the transcription start sites (TSS) of ER-Y537S specific up- and down-regulated genes and ER-Y537S enriched and depleted ER bound sites. Genomic distance analysis revealed that ER-Y537S enriched ER binding sites were located closer to ER-Y537S specific down-regulated genes (P-value = 2.28 x 10^-110^, Wilcoxon signed-rank test; Fig. 2C), while mutant-depleted ER binding sites were also significantly closer to mutant-specific down-regulated genes (P-value = 2.03 x 10^-71^, Wilcoxon signed-rank test; Fig. 2D). While the association between enriched binding sites and down-regulated genes appears to contradict ER’s primary role as an activating transcription factor, genes with ER-Y537S specific binding within 10 kb of their TSS are in fact enriched for genes up-regulated in ER-Y537S cells (P-value = 2.7 x 10^-12^, Wilcoxon signed-rank test) (Fig. 2C). The ER-Y537S distance analysis contrasts what was previously observed with ER-D538G, where around 40% of ER-D538G specific up-regulated genes were enriched for ER-D538G specific ER binding at a larger distance (within 100 kb) [22]. These results suggest the connection between ER binding and gene regulation may differ between mutations, with ER-Y537S typically acting at a shorter distance.

### ER-Y537S and ER-D538G alter chromatin accessibility

The findings that mutant ER affected gene expression and altered ER genomic binding in an allele-specific manner, led us to question whether the mutant receptors were able to impact chromatin accessibility. We performed the Assay for Transposase-Accessible Chromatin with high-throughput sequencing (ATAC-seq) in wildtype and ER-Y537S clones cultured in hormone depleted media and stimulated with 10nM E2 or DMSO for 1 hour. We then compared ER-Y537S data to our previously published wildtype and ER-D538G experiments [22] to identify how both mutations changed the chromatin landscape. When compared to wildtype cells, ATAC-seq analysis identified that ER-Y537S and ER-D538G altered chromatin accessibility at 5101 shared regions in the genome, with increased accessibility at 3768 loci and decreased accessibility at 1333 loci (Fig. 3A). In addition, ER-Y537S altered chromatin accessibility at an additional 210 (increased- 175, decreased- 35) regions and ER-D538G altered accessibility at an additional 861 (increased- 671, decreased- 190) regions. The differences in chromatin accessibility were largely shared between mutations, with a smaller role for mutation-specific chromatin accessibility patterns mediated by the different mutations, unlike the observed gene expression patterns.

**Figure 3.**
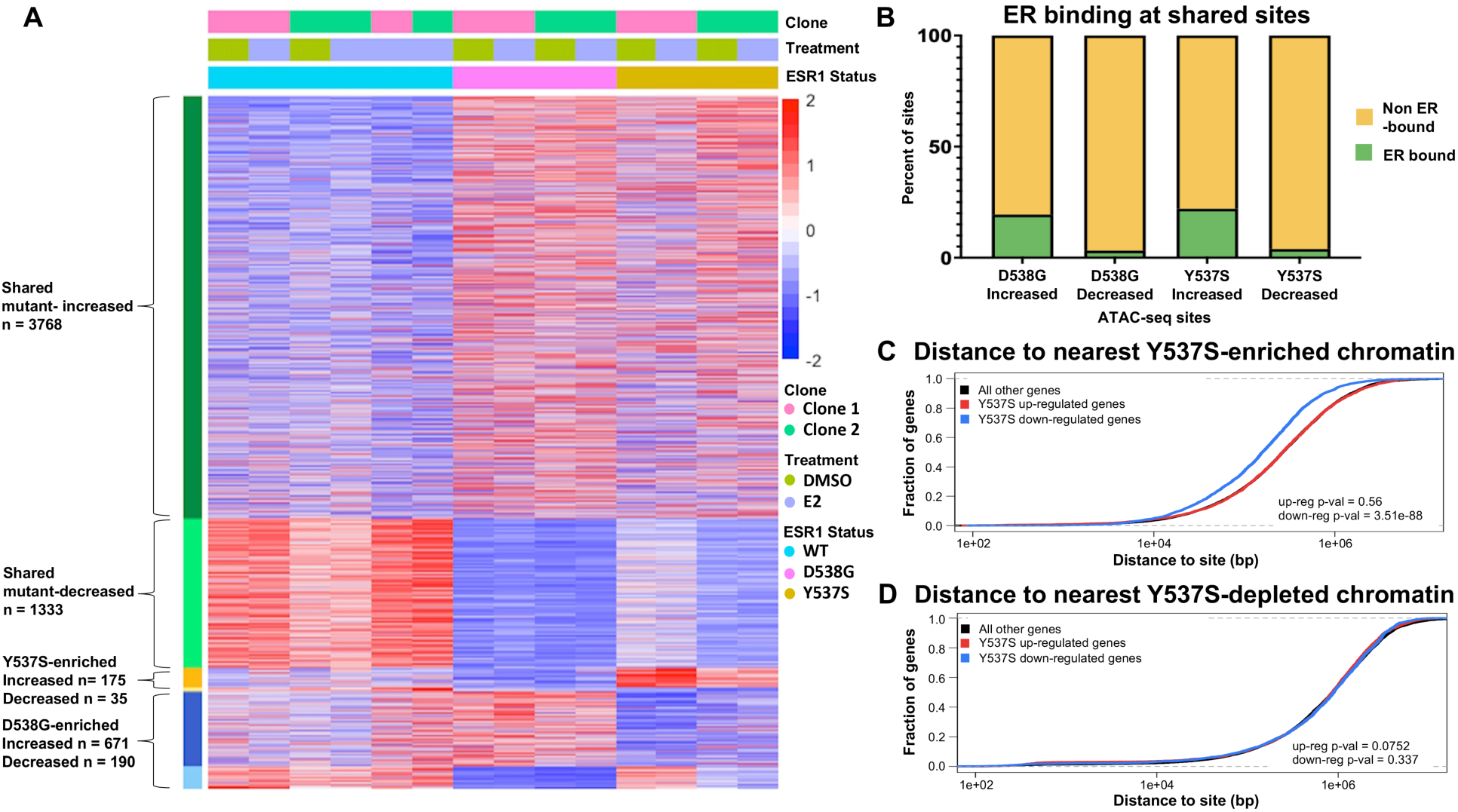
Mutant ER alters chromatin accessibility at shared and mutant-specific loci. (A) Heatmap displays ATAC-seq signal at shared mutant-enriched and shared mutant-depleted regions as well as ER-Y537S and ER-D538G enriched and depleted regions. (B) Bar graphs show the percent of differential chromatin sites that overlap ER genomic bound sites. Cumulative distribution graphs show distance from the transcription start sites of ER-Y537S specific genes to regions of ER-Y537S mutant-enriched (C) and depleted (D) accessible chromatin.

To better understand the role that ER binding could be playing in differential chromatin accessibility, we overlapped the ATAC-seq regions with ER bound regions from ChIP-seq experiments. Very few regions with decreased accessibility exhibited ER binding, while approximately 20% of ER-D538G and ER-Y537S enriched chromatin was associated with ER binding, suggesting that mutant ER may be directly contributing to increased accessibility at a minority of sites (Fig. 3B). Despite relatively few differential ATAC-seq sites exhibiting ER genomic binding, the allele-specific patterns of chromatin accessibility and ER genomic binding were consistent (Supplemental Fig. 3B). The overall pattern indicates that many of the changes to chromatin accessibility are not mediated by the mutant receptors through direct binding. Motif analysis in the non-ER associated regions implicated transcription factors AP-1 (P-value = 3.8 x 10^-44^, MEME) and CCCTC-Binding Factor (CTCF) (P-value = 7.1 x 10^-154^, MEME), suggesting other factors may mediate differential chromatin accessibility patterns. We next investigated how gene expression was related to nearby differential chromatin. Unexpectedly, 40% of ER-Y537S down-regulated genes were located within 100kb of ER-Y537S enriched chromatin (Fig. 3C). However, no significant enrichment of genes nearby ER-Y537S depleted chromatin was observed (Fig. 3D). Overall, our gene regulation results in endometrial cancer cells indicate that ER-Y537S causes larger changes in gene expression that are similar to constant ER activity, that ER-Y537S and ER-D538G have differences in their genomic binding, and that both mutations cause large scale chromatin accessibility changes that are mostly shared.

### ER-Y537S and ER-D538G do not alter proliferation of endometrial cancer cells in culture

The large number of genes differentially regulated by ER-Y537S and ER-D538G in addition to the enriched molecular and cellular functions, led us to investigate phenotypes being altered in ER mutant endometrial cancer cells. We first tested proliferation by culturing wildtype and ER-Y537S cells in full media and hormone depleted media for 72 hours and monitored their growth rates with live cell imaging via the IncuCyte Zoom platform. There were no observed differences in proliferation, with ER-Y537S cells growing at similar rates as wildtype cells in both media conditions. Consistent with findings from ER-D538G proliferation studies in Ishikawa cells, ER mutations do not confer a growth advantage *in vitro* [22] (Supplemental Fig. 4A).

### ER-Y537S and ER-D538G enhance migration and invasion of endometrial cancer cells

The downregulation of actins (Supplemental Fig. 4B) and cadherins (Supplemental Fig. 4C), which is typically associated with increased movement, led us to investigate whether ER-Y537S affected migration in our models. To test this hypothesis, we performed a wound healing assay, seeding wildtype and ER-Y537S cells in full serum media or hormone depleted media. ER-Y537S cells migrated 45% faster than wildtype cells in full serum media (Fig. 4A; P-value <0.0001, unpaired t-test) (examples in Fig. 4B) and 67% faster than wildtype cells in hormone depleted media (Fig. 4C; P-value <0.0001, unpaired t-test) (examples in Fig. 4D). The trend of increased migration of ER-Y537S models in endometrial cancer is similar to our previous findings with ER-D538G cell lines (Fig. 4A-4D), suggesting that ER mutant endometrial cancer cells have the potential for increased migration, one of the hallmarks of more aggressive disease [49, 50].

**Figure 4.**
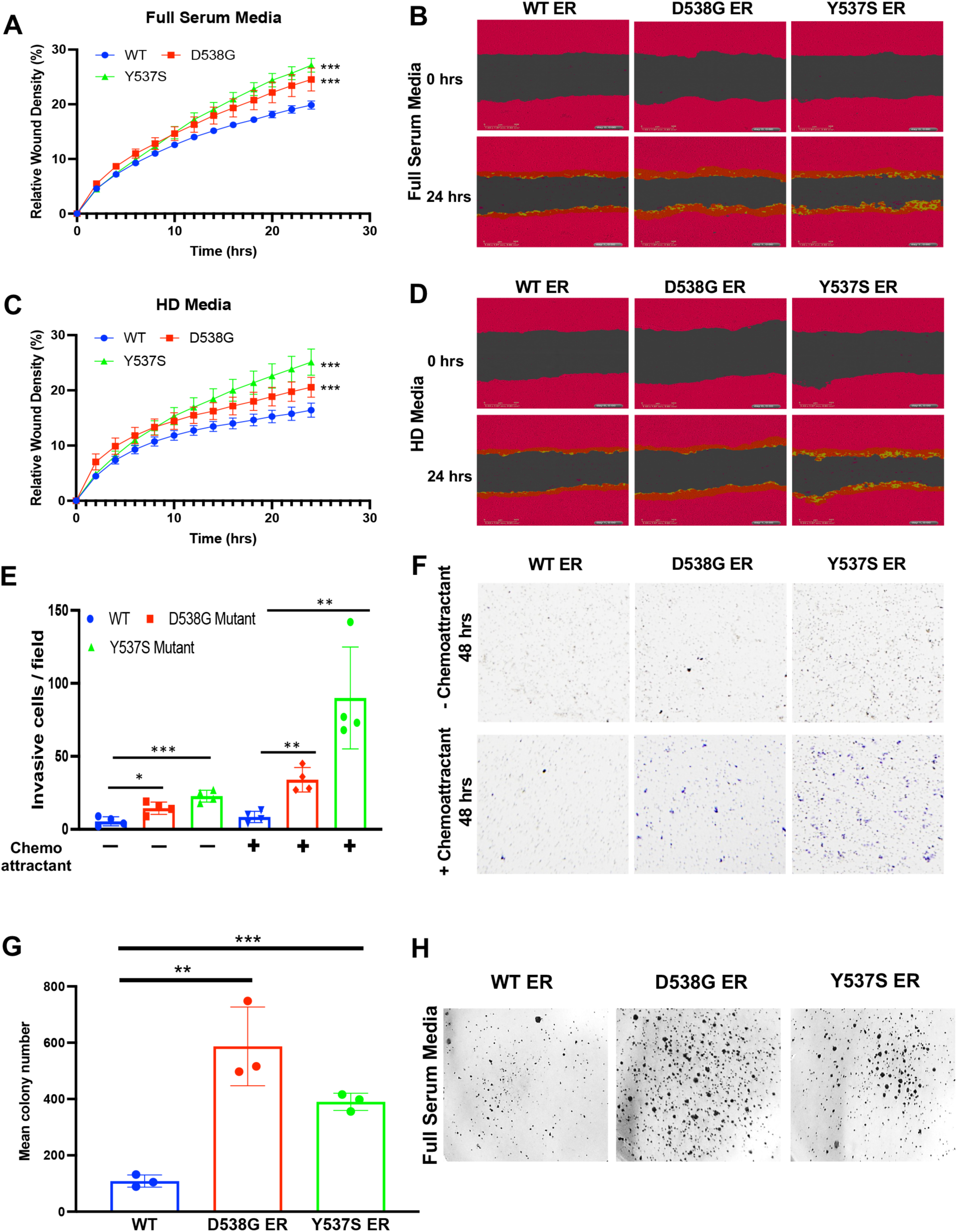
Mutant ER promotes migration, invasion and increased anchorage-independent growth in endometrial cancer. Increased rates of migration were observed for ER-D538G and ER-Y537S cell lines as measured by the relative scratch wound density in full serum media. (A) and hormone depleted (HD) media (C) are shown. Indicated p-values are relative to wildtype. Images show cell monolayer at 0 hours and 24 hours post wound scratch in full serum media (B) and HD media (D) for wildtype, ER-D538G, and ER-Y537S cell lines. (E) Bar graph shows the number of ER-WT, ER-D538G, and ER-Y537S cells per field that have invaded the extracellular matrix (matrigel) in the absence and presence of FBS. (F) Representative images show invasive cells in the absence and presence of FBS at 48 hours. The relative number of invasive cells were calculated using ImageJ software. (G) Graph displays mean number of ER-WT, ER-D538G, and ER-Y537S colonies formed in soft agar after 27 days in full serum media. (H) Representative images show relative number of colonies formed in soft agar after 27 days. * p < 0.01; ** p < 0.001; *** p < 0.0005.

Studies have suggested that nutrient deprivation is a major feature of the tumor microenvironment and under short periods of limited nutrients cells lose their migratory and proliferative potential [51]. To further elucidate this phenotype in ER mutant models, we tested the ability of wildtype, ER-Y537S, and ER-D538G cells to migrate in the context of nutrient deprivation. ER-D538G cells migrated 35% faster than wildtype controls and ER-Y537S mutant cells migrating 58% faster than wildtype cells in nutrient deprived conditions (Supplemental Fig. 4D). These results not only confirm the migratory phenotype in ER mutant endometrial cancer cells but also suggest that ER-Y537S mutant cells are significantly more migratory and have the potential to be more aggressive than ER-D538G endometrial cancer cells.

To follow up on the observation that ER mutant cells were significantly more migratory, we analyzed the ability of the mutations to promote cell invasion, via a transwell invasion assay. At 48 hours, we found ER mutant cells invaded the matrigel significantly more than wildtype cells in the absence and presence of chemoattractant (Fig. 4E, examples in Fig. 4F). Corroborating our migration findings, both mutations were more invasive than wildtype cells, but ER-Y537S cells were more invasive than ER-D538G mutant cells. Taken together these experiments demonstrate that ER mutations play an important role in promoting increased migration and invasion in endometrial cancer cells and indicate potentially more aggressive tumors than their wildtype counterparts, especially in the context of ER-Y537S. The ER mutant associated phenotypes are consistent with enriched cellular functions in mutant regulated genes (Supplemental Fig. 1C, D).

### ER-Y537S and ER-D538G promote colony formation in soft agar

The increased migration and invasion of ER mutant cells suggests that both mutations can impact cancer related phenotypes. One phenotype that is often a feature of transformation in cancer is the ability of cells to proliferate in anchorage-independent conditions. To functionally test this phenotype, wildtype, ER-Y537S, and ER-D538G cells were grown in 0.5% agar for up to 27 days and monitored for colony formation over time. We observed that ER-Y537S and ER-D538G cells formed significantly more colonies than wildtype controls (Fig. 4G) (examples in Fig. 4H). ER-D538G cells had a selective advantage in anchorage-independent conditions, forming an increased number of colonies in the assay when compared to ER-Y537S cell lines. Although the majority of the phenotypic experiments suggested increased activity of ER-Y537S, this unexpected finding from the anchorage independent experiments suggest that ER-D538G also has the potential to make cells more aggressive and possibly metastatic in endometrial cancer.

### ER-Y537S promotes tumor growth *in vivo*

Despite the inability of ER mutants to provide a proliferative advantage in cell culture experiments, we hypothesized that the increased activity of the mutant receptors could provide a growth advantage *in vivo*, as observed in breast cancer models [5, 6, 12, 14]. To test this hypothesis, wildtype, ER-Y537S, or ER-D538G cells were injected into the mammary fat pad of ovarectomized immunodeficient mice in the presence or absence of E2 pellets. Consistent with *in vitro* experiments we observed that at 35 days post cell implantation, all mice that received estrogen supplementation showed no significant differences in tumor growth rates (Fig. 5A). When comparing mice that did not receive estrogen supplementation, tumor growth was significantly increased in ER-Y537S models compared to ER-D538G and wildtype models (Fig. 5B). This data suggests that despite the elevated activity of both mutations, only ER-Y537S was able to compensate for the lack of estrogens and promote tumor growth *in vivo*.

**Figure 5.**
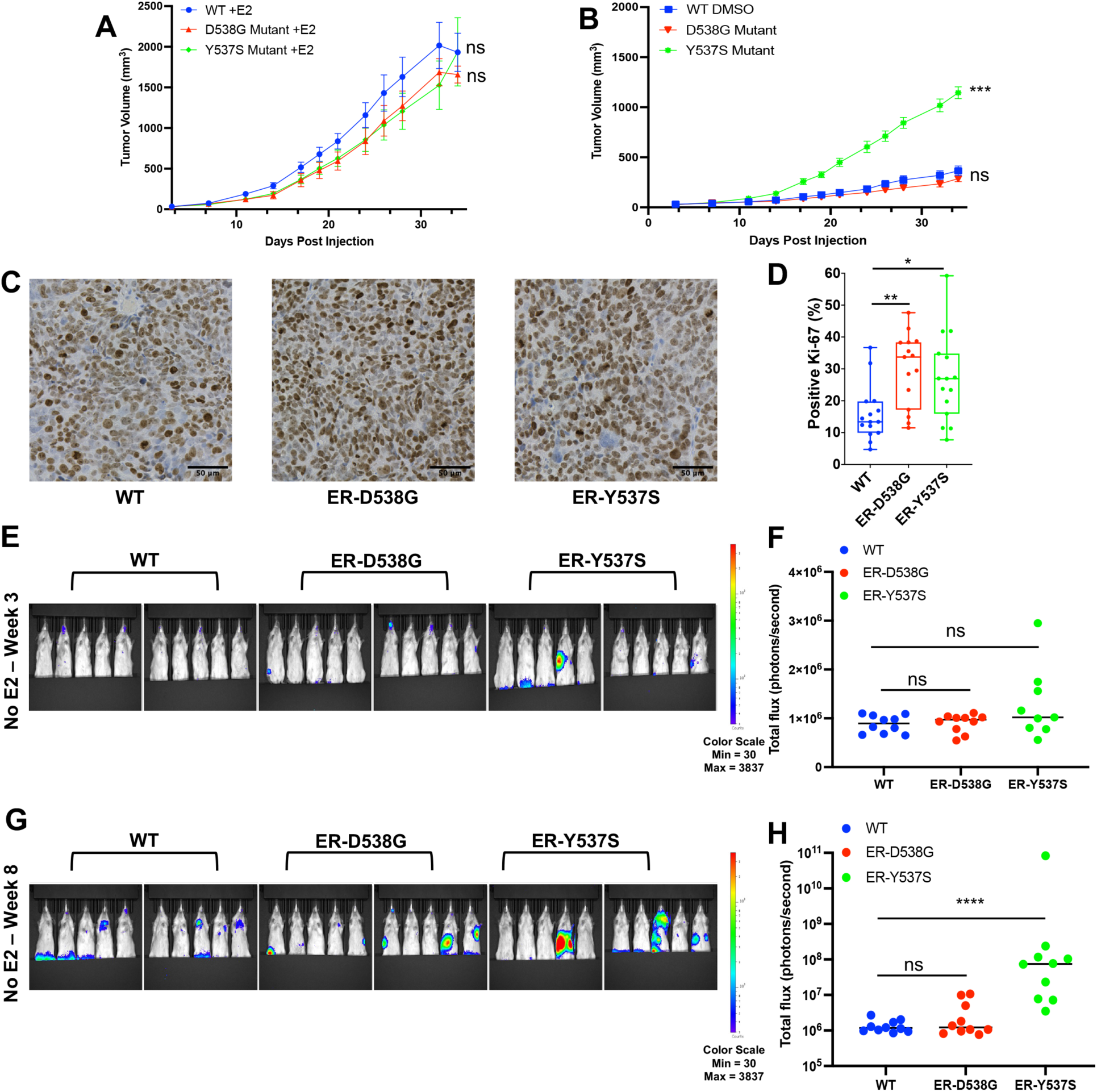
ER-Y537S drives tumor formation *in vivo* and promotes metastasis. Tumor growth is shown after ER-WT, ER-D538G, and ER-Y537S cells were injected into ovarectomized and immunocompromised mice with (A) and without (B) E2 supplementation. (C) Representative fields of Ki-67 staining for WT, ER-D537G, and ER-Y537S xenograft tumors are shown. Scale bar represents 50 μm. (D) Box and whisker plot shows Ki-67 positivity across 15 individual fields for WT, ER-D538G, and ER-Y537S xenograft tumors. P-values calculated by ANOVA with Tukey’s post-hoc multiple comparisons test. Whole body IVIS images depict luminescence signal ovariectomized mice 3 weeks (E) and 8 weeks (G) post tail vein injection with wildtype, ER-D538G and ER-Y537S cells expressing luciferase. Quantification of luminescence in whole animals is shown at week 3 (F) and week 8 (H). *P* values calculated via Mann-Whitney U test. * p < 0.05, ** p < 0.01; *** p < 0.0005; **** p < 0.0001; n.s. not significant.

To further assess the consequences of ER mutations *in vivo* and identify differences between the ER-Y537S and ER-D538G mutations, we performed RNA-seq on xenografts at the end of the experiment without E2 supplementation. We identified 4544 genes differentially up- (2241) and down-regulated (2303) by both the ER-Y537S and ER-D538G xenografts (Supplemental Fig. 5A; for gene lists, see Supplemental Table 3). These shared differentially expressed genes were enriched for terms relating to proliferation, DNA replication, and cell cycle progression, suggesting that both mutations drive proliferation *in vivo* (Supplemental Fig. 5B). We observed the hallmark signatures relating to estrogen response as increased in the shared differentially expressed genes as well, suggesting that mutant ER retains estrogen independence *in vivo* (Supplemental Fig. 5C). We also identified 3140 genes differentially up- (1644) and down-regulated (1496) specifically in the ER-Y537S xenografts, while 2408 genes were differentially up- (1207) and down-regulated (1201) in the ER-D538G xenografts compared to wildtype tumors (Supplemental Fig. 5A; for gene lists, see Supplemental Table 3). Genes upregulated by the ER-Y537S xenografts were enriched for catabolic processes, oxidative phosphorylation, and hypoxia, whereas genes differentially expressed in the ER-D538G xenografts were enriched for transcriptional regulation, DNA repair and replication, and the cell cycle (Supplemental Fig. 5B-C).

Despite the ER-D538G tumors being of similar size to wildtype tumors, gene expression suggests that both ER-D538G and ER-Y537S tumors are more proliferative than wildtype tumors. Therefore, we assessed the proliferative marker Ki-67 in the xenograft tumors to determine proliferation rates (Fig. 5C). We found that both the ER-Y537S and ER-D538G xenografts had significantly higher Ki-67 positivity than wildtype xenografts, consistent with RNA-seq findings (Fig. 5D). Furthermore, H&E staining of the xenografts showed that the wildtype and ER-Y537S tumors have large regions of necrosis, consistent with increased hypoxia in ER-Y537S, compared to the ER-D538G tumors (Supplemental Fig. 5D). This may explain how the ER-D538G tumors are more proliferative than wildtype tumors despite having similar tumor volume, as much of the wildtype tumors are necrotic. These findings also indicate that ER-D538G tumors may have been poised to increase growth at the end of the experiment, which is consistent with breast cancer ER-D538G tumors taking longer to grow *in vivo* [14].

We next examined the role of ER mutants in promoting metastases in immunocompromised mice. Wildtype, ER-Y537S and ER-D538G cells expressing luciferase were injected through the tail vein into ovariectomized mice with no estrogen supplementation. Mice were IVIS imaged weekly for luminescence and monitored for metastatic burden over time. When comparing an earlier week 3 timepoint (Fig. 5E-F) to the later week 8 timepoint (Fig. 5G-H), we observed the whole mouse metastatic burden significantly increased over time, in mice injected with ER-Y537S cells compared to the other genotypes. ER-Y537S tumors were observed in the lungs and liver, with one ER-Y537S animal exhibiting a brain metastasis, an interesting event that is rarely observed in patients with endometrial carcinoma [52]. Consistent with the increased molecular and phenotypic activity, these results suggest ER-Y537S endometrial cancer cells are significantly more aggressive and can metastasize more readily.

### RIME identifies ER associated cofactors and potential drug targets in endometrial cancer

The increased estrogen-independent phenotypes of ER mutant endometrial cancer cells are consistent with more aggressive disease and indicate a need for tailored therapeutic options. One possible approach towards treating patients with ER mutant tumors is to target cofactors that are necessary for ER’s gene regulatory function. ER is known to recruit co-activators necessary for target gene expression and we hypothesized that blocking ER cofactors, which could be endometrial cancer-specific, is an effective strategy for blocking mutant ER activity. By exploiting ER’s cofactors as therapeutic targets for ER mutant or estrogen-driven endometrial cancer, we could identify a novel strategy for the treatment of this disease.

Despite the major oncogenic role that ER plays in endometrial cancer, ER coregulatory proteins in this disease remain relatively unknown. In breast cancer, several regulatory proteins that help ER regulate gene expression have been found, including Forkhead box A1 (FOXA1), GATA Binding Protein 3 (GATA3), Growth Regulating Estrogen Receptor Binding 1 (GREB1), and Nuclear Receptor Coactivator 3 (NCOA3/SRC-3) [42, 53–55]. To gain insight into endometrial cancer ER associated cofactors, we performed RIME in Ishikawa parental cells and compared the data to published breast cancer ER RIME datasets. This analysis identified some established ER interactors such as NCOA3 and Histone Acetyltransferase P300 (EP300) (Fig. 6A); however, most ER co-occurring factors were not shared between breast and endometrial cancer cells, suggesting that we have identified unique endometrial cancer-specific ER associated proteins (Supplemental Fig. 6A, Supplemental Table 5). To analyze proteins that co-occur with mutant ER, we performed RIME in the ER-Y537S and ER-D538G cells. We found that the majority of wildtype ER co-occurring factors could be observed in the ER mutant cells (86% for ER-Y537S and 80% for ER-D538G; Supplemental Table 5). While no novel significant cofactors were identified for ER-Y537S or ER-D538G, some cofactors exhibited increased RIME signal in the ER mutant cells (14% for ER-Y537S and 6% for ER-D538G; Supplemental Table 5). These results indicate that mutant ER utilizes similar coregulatory proteins as wildtype ER while maintaining cancer type specificity.

**Figure 6.**
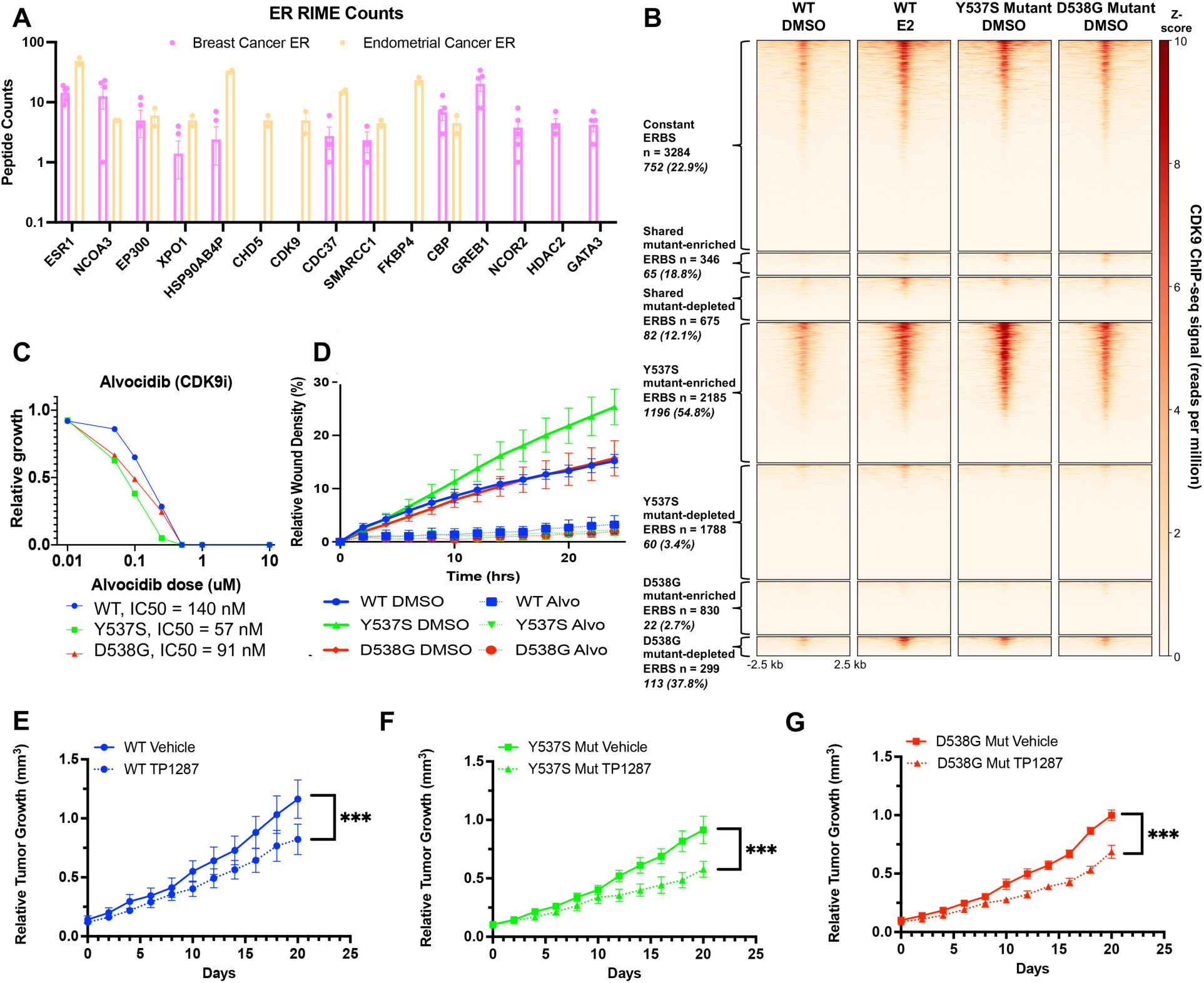
Exploring novel therapeutic targets in ER active and mutant endometrial cancer. (A) RIME peptide count data shows ER co-associating proteins in breast cancer and endometrial cancer cell lines. (B) Heatmap shows CDK9 binding patterns at ER bound sites (ERBS) in wildtype, ER-Y537S, and ER-D538G cell lines. The number of ERBS in each category is given, with the overlapping number and percent of CDK9 bound sites shown in italics. (C) Relative growth is shown for wildtype, ER-Y537S, and ER-D538G cell lines after treatment with Alvocidib (CDK9 inhibitor) for 72 hours. (D) Reduced rates of migration are observed after CDK9 inhibition, as measured by the relative wound scratch density of wildtype, ER-D538G, and ER-Y537S cell lines in full serum media. Inhibition of CDK9 reduces tumor burden in wildtype (E), ER-D538G (F), and ER-Y537S (G) xenograft models. Mice were ovariectomized and received estrogen supplementation. Untreated controls n = 10 mice/group, treated mice n = 8 mice/group. *** p < 0.0001.

Factors that interact with ER, as identified by RIME, could be explored as therapeutic targets in ER active and ER mutant endometrial cancer. Based on the list of co-occurring factors, we prioritized two cofactors that co-occurred with wildtype and mutant ER (Supplemental Fig. 6B-C) and had available targeting agents: NCOA3, also known as Steroid Receptor Coactivator 3 (SRC-3), a protein that has been shown to interact with ER in breast cancer models [18], and Cyclin Dependent Kinase 9 (CDK9), an endometrial cancer-specific co-occurring ER protein that is a member of the P-TEFb complex [56]. As CDK9 is a novel ER co-occurring factor in endometrial cancer cells with increased RIME signal in ER mutant cells, we first sought to describe the genomic relationship between ER and CDK9.

Western blot analysis identified that CDK9 protein levels were unchanged in ER mutant cells (Supplemental Fig. 6D). To determine the genomic binding relationship between ER and CDK9, we performed ChIP-seq for CDK9 in wildtype, ER-D538G, and ER-Y537S cells without E2 supplementation, along with wildtype cells with E2 supplementation. Assessing CDK9 binding alone, we found 48 regions (44 up, 4 down) that are consistently differentially bound by CDK9 in wildtype cells treated with E2, ER-Y537S, and ER-D538G compared to wildtype cells without E2 treatment (Supplemental Fig. 6E). In addition, we identified 317 regions (190 up, 127 down) that exhibited differential binding in both ER mutant lines compared to wildtype (Supplemental Fig. 6E). We further identified genotype-specific CDK9 differential binding when comparing to wildtype cells without E2 treatment, with 455 regions (449 up, 6 down) specific to wildtype cells treated with E2, 1,098 regions (826 up, 272 down) specific to ER-Y537S cells, and 844 regions (369 up, 475 down) specific to ER-D538G expressing cells (Supplemental Fig. 6E). We also found a high level of overlap between CDK9 and ER genomic binding. CDK9 was bound to several regions that were also bound by ER (Fig. 6B) and CDK9 ChIP-seq signal at ER bound sites was correlated with patterns in ER binding (Supplemental Fig. 6F). The high occupancy of CDK9 at ER bound sites and binding signal that changes coordinately with changes in mutant ER binding further support the connection between ER and CDK9, which was identified by RIME, and exploration of CDK9 as a novel therapeutic target.

### Novel therapeutic targets in ER mutant endometrial cancer

The RIME analysis revealed multiple endometrial cancer ER co-occurring factors that could be explored as therapeutic targets in ER active and ER mutant endometrial cancer. We tested the functional consequences of SRC-3 and CDK9 inhibition in endometrial cancer cell lines *in vitro* by using small molecule inhibitors SI-2 and Alvocidib, respectively. We first examined the effects of SI-2 and Alvocidib on proliferation in Ishikawa wildtype and ER mutant cells over a range of drug doses. Treatment with SI-2 and Alvocidib as single agents resulted in dose-dependent decreases in proliferation in wildtype and ER mutant cell lines. SI-2 displayed similar growth inhibition across wildtype and ER mutant cells (Supplemental Fig. 7A). For Alvocidib, ER-Y537S and ER-D538G cell lines were more sensitive to CDK9 inhibition compared to wildtype controls, with the ER-Y537S lines displaying the highest sensitivity in these experiments (Fig. 6C). We next tested the ability of both drugs to inhibit the increased migration observed in ER mutant cell lines. Treatment with SI-2 decreased migration by 31% in ER-D538G cell lines and by 36% in ER-Y537S lines (Supplemental Fig. 7B). Alvocidib treatment resulted in more than 90% reduction in migration in both ER mutant cell lines (ER-D538G – 91%; ER-Y537S – 94%; Fig. 6D), suggesting that targeting CDK9 may be a particularly powerful way to diminish the ability of ER mutant endometrial cancer cells to both migrate and grow.

Based on the positive results of *in vitro* treatment, we tested the *in vivo* efficacy of SRC-3 and CDK9 inhibition, creating xenograft models with wildtype, ER-Y537S, and ER-D538G cells implanted in the flank of ovariectomized mice with E2 supplementation. Once tumors reached 100 – 200 cubic millimeters, mice were randomized and treated with either SI-2, TP-1287 (an oral phosphate prodrug of Alvocidib with enhanced pharmacokinetics), or vehicle control, and we monitored their response to treatment over time. SI-2 treated mice showed a slight reduction in tumor growth 20 days post implantation compared to vehicle controls, regardless of ER genotype (Supplemental Fig. 7C-E). However, this trend did not reach significance in our cohorts. This suggests SRC-3 inhibition as a monotherapy may not be sufficient to reduce tumor growth. When comparing TP-1287 treated mice to controls, CDK9 inhibition significantly slowed tumor growth over time in all genotypes, with a greater reduction in both ER mutant models (Fig. 6E-G). Histopathological analysis of hematoxylin and eosin (H&E) stained tumor sections from each treatment group revealed that although tumors from each genotype grew at different rates, they retained an endometrioid histology regardless of genotypes and treatment (Supplemental Fig. 7F). There were differences in the degree of necrosis, with ER-Y537S SI-2 treated tumors showing less extensive necrosis than wildtype and ER-D538G treated tumors. Although there was not a complete response to CDK9 inhibition, the data suggests that CDK9 inhibition is a promising therapeutic tool that could be explored further.

## DISCUSSION

Despite the prevalence and implied estrogen-independent activity, little is known about ER mutant function and contribution to disease progression in endometrial cancer. In this study, we interrogated the common ER-Y537S and ER-D538G mutations, systematically profiling their molecular and phenotypic consequences, and identified potential therapeutics for the treatment of ER mutant disease. RNA-seq experiments performed *in vitro* confirmed the ligand independent activity of mutant ER in addition to allele-specific transcriptional regulation of thousands of genes. These results are similar to breast cancer studies that show allele-specific transcriptional regulation by mutant ER, although the genes that are impacted differ between cancer types. The molecular basis for the differences between these adjacent mutations is unclear, but our analysis revealed that ER-Y537S gene expression effects were more similar to sustained estrogen signaling in the wildtype context compared to ER-D538G mediated expression changes. This indicates that ER-Y537S could represent a more active form of ER while ER-D538G might possess additional properties. Mutation-specific expression changes showed downregulation of cellular functions associated with movement in addition to an association with genes differentially expressed in ER mutant endometrial cancer patient tumors and with genes whose expression is correlated with worse endometrial cancer patient outcomes. These findings suggest that ER-Y537S and ER-D538G regulate similar processes in endometrial cancer that are consistent with more aggressive disease.

To determine how mutant ER impacted gene expression, we analyzed genomic binding of mutant ER using ChIP-seq. We found that both mutant receptors significantly reprogrammed ER binding site selection, however ER-Y537S showed increased binding capacity. Additionally, ER-Y537S utilized some promoter-mediated interactions, while ER-D538G displayed mostly enhancer-mediated regulation in endometrial cancer. This demonstrates ER mutations dictate not only where the mutant receptors bind in the genome but how this binding relates to gene expression. We questioned how other genomic alterations could contribute to the allele-specific activity in our models and assessed changes to the chromatin landscape via ATAC-seq. The majority of changes to the chromatin landscape were shared between mutations. Mutant ER could be playing a direct role in approximately 20% of ER-Y537S and ER-D538G enriched chromatin accessible regions; however, a large portion of differential regions did not exhibit direct ER binding. This data suggests that some mutation-specific transcriptional and genomic differences we have identified can be attributed to differential chromatin accessibility with other transcription factors, such as AP-1 and CTCF, potentially facilitating some of the observed expression differences. In agreement with this concept, our previous studies in breast cancer found that CTCF is important in mediating the mutant ER gene expression program [13].

To understand how the molecular findings from ER mutant studies impacted phenotypes, we functionally interrogated cancer related cellular processes. ER mutations did not affect cell growth but promoted increased migration and invasion. ER-Y537S was significantly more migratory and invasive than ER-D538G, consistent with the increased gene expression differences. The increased ER-Y537S activity may suggest more aggressive endometrial tumors, which parallel findings from the BOLERO2 breast cancer clinical trial, which concluded that ER-Y537S patients had poorer clinical outcomes compared to ER-D538G tumors [57–59]. ER mutations also significantly increased colony formation in soft agar compared to wildtype cells. In contrast to migration and invasion, ER-D538G formed more colonies in anchorage-independent conditions, another finding consistent with breast cancer ER-D538G MCF7 models [12]. These unexpected results suggest an ER-D538G mediated role in metastasis, which may be shared between cancer types.

Given the increased molecular activity and ability of ER mutants to adapt and survive, we investigated their ability to promote tumor formation *in vivo*. Estrogen treated mice had similar growth rates, suggesting that co-expression of the wildtype allele has the ability to mask the estrogen-independent activity of the mutant; another finding shared with breast cancer models [17]. In the absence of estrogen supplementation, tumor growth was significantly increased in ER-Y537S implanted mice compared to ER-D538G and wildtype implanted animals. These results suggest an ER-Y537S mediated estrogen-independent mechanism, resulting in faster growing and more aggressive tumors *in vivo*. Consistent with these results, the *in vivo* metastasis study confirmed ER-Y537S was significantly more aggressive, with ER-Y537S animals showing metastases in several organ sites. The increased growth and metastatic potential of ER-Y537S tumors may be related to its association with constant ER activity, since we’ve shown previously that Ishikawa cells depend on E2 for growth *in vivo* and that having fully functional estrogen signaling is important for this E2 stimulated growth [46]. Despite the reduced ability of ER-D538G to form tumors *in vivo* in the absence of E2 during our time course, the increased activity of the mutant receptor *in vitro* and its more unique gene expression program suggests that ER-D538G tumors may need longer latency to adapt in animal models. This concept is supported by our *in vivo* gene expression analysis of ER mutant models, which uncovered elevated expression of cell proliferation genes compared to wildtype tumors, a finding that was confirmed by Ki-67 staining. The idea that ER-D538G is estrogen independent *in vivo* but with a longer latency than ER-Y537S is supported in breast cancer models where ER-D538G takes longer to grow *in vivo* [14].

We hypothesized that an understanding of allele-specific regulation by mutant ER could lead to the identification of targeted therapies to treat patients with ER mutant disease. Currently, new approaches are being used in clinical breast cancer models, with newer generation SERMs, such as bazedoxifene and lasofoxifene, being tested as single agents or in combination with Palbociclib, a CDK 4/6 inhibitor, against ER mutant disease [17, 60]. Additionally, newer generation SERDs such as giredestrant and elacestrant, which have better bioavailability than older compounds like fulvestrant, have also reached Phase 2 clinical trials [61]. Other treatment strategies include profiling coactivating proteins that bind to the mutant ER in an effort to identify novel therapies [18]. New strategies like this approach are promising as multiple studies have shown that ER mutations are able to bypass hormone therapies [10, 12, 14, 15]. Adopting a cofactor-focused approach in endometrial cancer, we utilized RIME to discover ER’s co-occurring factors and identify targeted therapies. We identified ER co-occurring proteins in endometrial cancer, some of which are shared with breast cancer, but many that are cancer type-specific. These findings are significant, as ER coregulatory proteins have remained under explored in the context of endometrial cancer and the candidate factors discovered by RIME lay the foundation for future studies on ER’s activity in endometrial cancer.

We investigated the potential of inhibiting two ER associated proteins, SRC-3 and CDK9, in the treatment of ER active and mutant endometrial cancer because of the availability of inhibiting compounds and consistent RIME signal in both ER wildtype and mutant endometrial cancer cells. The use of small molecule inhibitors resulted in decreased proliferation and migration in wildtype and ER mutant cells, with ER mutant lines exhibiting increased sensitivity to CDK9 inhibition. *In vivo* testing of the SRC-3 inhibitor revealed a slight decrease in tumor growth in all genotypes, although the single agent treatment was not sufficient for significant tumor regression. We used the SRC-3 inhibitor SI-2 in our studies; however, SI-12 is an improved version that was recently published and may have more efficacy in these models [62]. The *in vivo* experiments showed significant but partial tumor growth reduction following CDK9 inhibition in wildtype and ER mutant models. CDK9 is known to interact with the bromodomain protein BRD4 at ER bound enhancers [63], suggesting that combination treatments with BET inhibitors may reduce tumor growth further. It is also possible that other treatments may be effective in combination with CDK9 inhibition. In summary, we modeled the most clinically prevalent ER mutations in endometrial cancer and systematically compared their molecular and phenotypic effects. The findings point to differences between individual mutations, despite their regulation of similar processes. The exploration of ER’s interacting proteins revealed candidates for therapeutic targeting that have not been previously explored in the treatment of ER mutant endometrial cancer and could represent new treatment strategies.

## Acknowledgements

This research was supported by a Department of Defense Breast Cancer Research Program Breakthrough Award to Jason Gertz (W81XWH-16-1-0421) and the Huntsman Cancer Institute. Research reported in this publication utilized the Preclinical Research Shared Resource and the High-Throughput Genomics Shared Resource at Huntsman Cancer Institute at the University of Utah and was supported by the National Cancer Institute of the National Institutes of Health under Award Number P30CA042014. We thank the Oregon Health and Science University Proteomic Shared Resource Core, Bryan Welm, Ben Spike, Thanh Parks, Margit Janat-Amsbury, and K-T Varley as well as Gertz and Varley laboratory members for their helpful comments on the study and the manuscript. We also thank Sumitomo Pharma Oncology for providing us with TP-1287 for the *in vivo* drug studies.

