## Supplemental Figures for "Allele-specific gene regulation, phenotypes, and therapeutic vulnerabilities in estrogen receptor alpha mutant endometrial cancer"

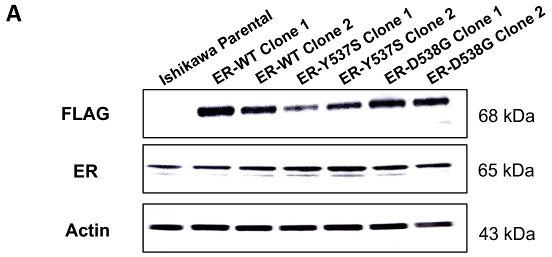


B

| Molecular and Cellular Functions [Y537S genes] | Direction | p-value |
|---|---|---|
| mRNA binding | ↑ | 5.2E-10 |
| Protein serine/threonine kinase activity | ↑ | 1.2E-6 |
| Actin binding | ↓ | 4.6E-4 |
| Actin filament binding | ↓ | 7.2E-4 |

C

| Molecular and Cellular Functions [D538G genes] | Direction | p-value |
|---|---|---|
| Phospholipase activity | ↑ | 2.6E-3 |
| Histone demethylase activity (H3-K9 specific) | ↑ | 4.6E-3 |
| Actin binding | ↓ | 1.6E-5 |
| Cadherin binding | ↓ | 2.7E-4 |

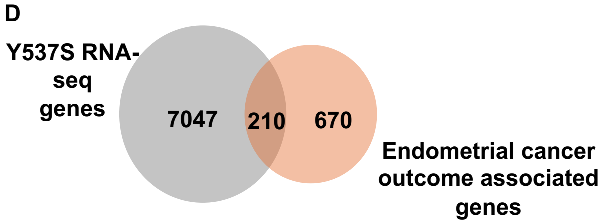


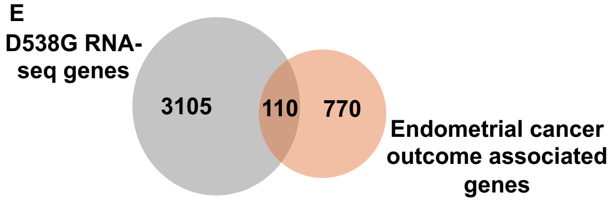


**Supplemental Figure 1. Validation and gene expression analysis of ER mutant models (related to Figure 1).** (A) Immunoblotting in Ishikawa parental, two each of heterozygous FLAG-tagged ER-WT, ER-Y537S, and ER-D538G cell lines show protein levels of FLAG epitope tagged ER, total ER, and Actin. Enrichr analysis highlighting molecular and cellular functions enriched in ER-Y537S (B) and ER-D538G (C) mutant-specific up and downregulated genes. Venn diagrams indicate the overlap between ER-Y537S specific up- and down-regulated genes (D) and ER-D538G specific up- and down-regulated genes (E) and genes associated with endometrial cancer patient outcomes in TCGA studies.

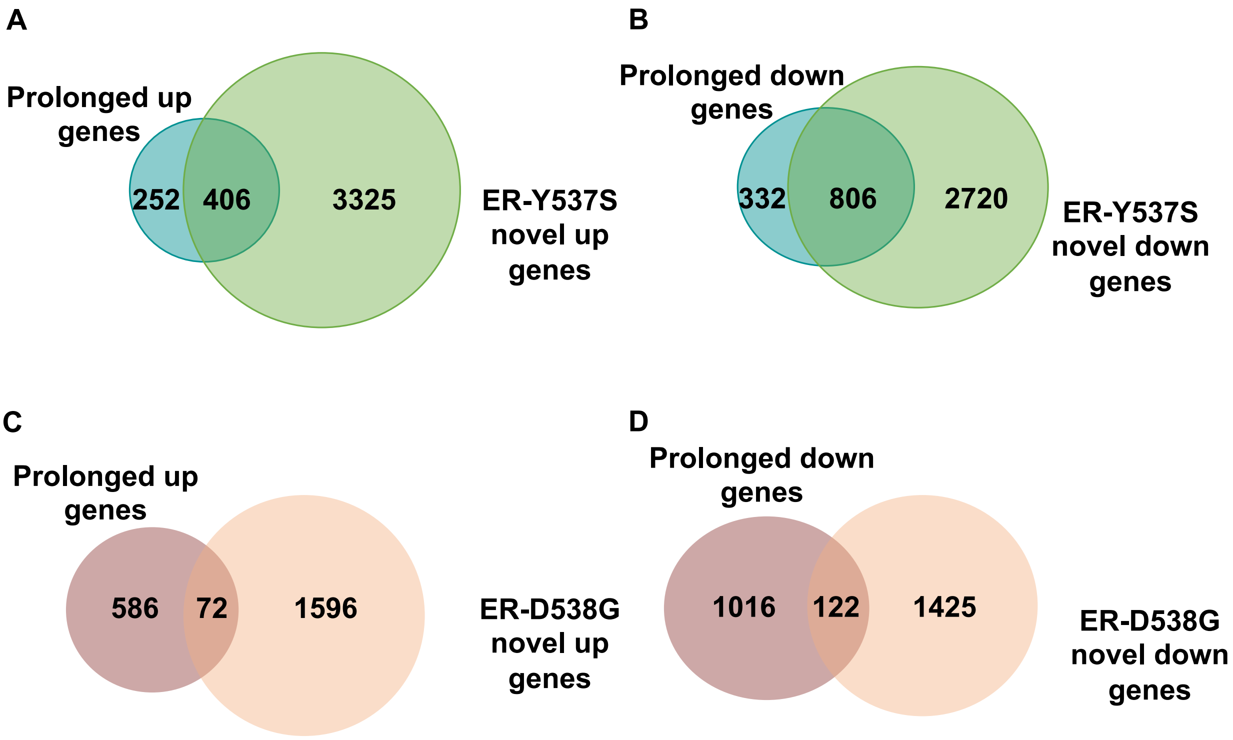


**Supplemental Figure 2. Comparison of mutant ER regulated genes and gene regulated by sustained estrogen signaling (related to Figure 1).** (A) Venn diagram shows the overlap between genes up-regulated by prolonged E2 and ER-Y537S specific up-regulated genes (4.36 fold enrichment, p-value $1.97e^{-180}$; hypergeometric test). (B) Venn diagram shows the overlap between genes down-regulated by prolonged E2 and ER-Y537S specific down-regulated genes (5.3 fold enrichment, p-value $1.67e^{-462}$; hypergeometric test). (C) Venn diagram shows the overlap between genes up-regulated by prolonged E2 and ER-D538G specific up-regulated genes (1.73 fold enrichment, p-value $4.2e^{-6}$; hypergeometric test). (D) Venn diagram shows the overlap between genes down-regulated by prolonged E2 and ER-D538G specific down-regulated genes (1.83 fold enrichment, p-value $6.79e^{-11}$; hypergeometric test).

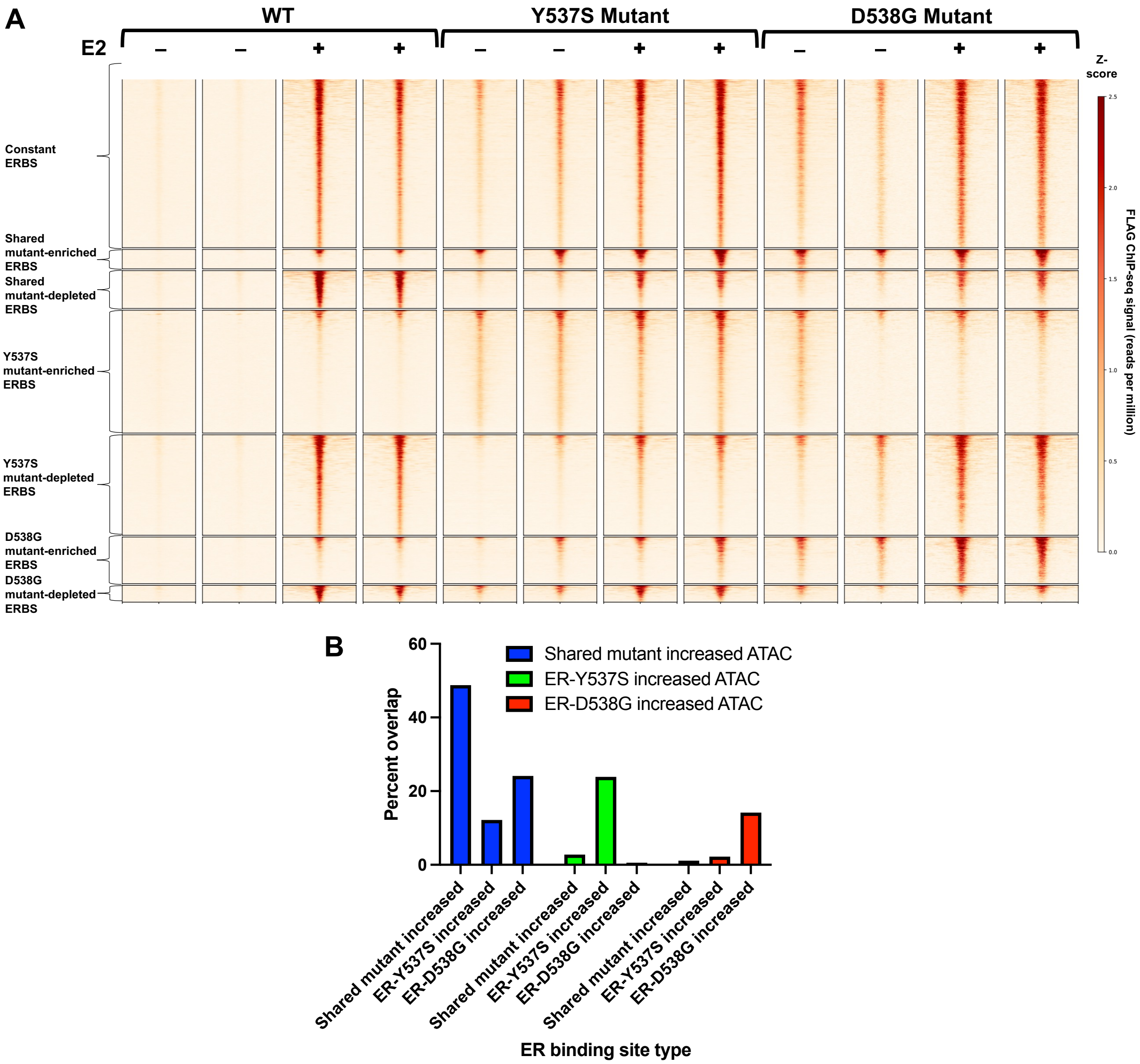


**Supplemental Figure 3. ER binding is altered by the Y537S and D538G mutations (related to Figures 2 and 3).** (A) Heatmaps display wildtype, ER-Y537S, and ER-D538G mutant binding in hormone depleted media following treatment with DMSO or E2 for one hour. ER binding sites include constant regions that are similar in wildtype and mutant lines (top panels), sites that are enriched or depleted and shared between ER-Y537S and ER-D538G cell lines (middle panels), and sites that are enriched or depleted in a mutant-specific manner (bottom panels). Each line is a an ERBS site, regions represent 2.5kb up- and down-stream of the peak summit. (B) Bar graph shows the overlap between mutant-enriched ATAC-seq sites and mutant-enriched ER bound sites. In each case, the denominator is the smaller of the two sets.

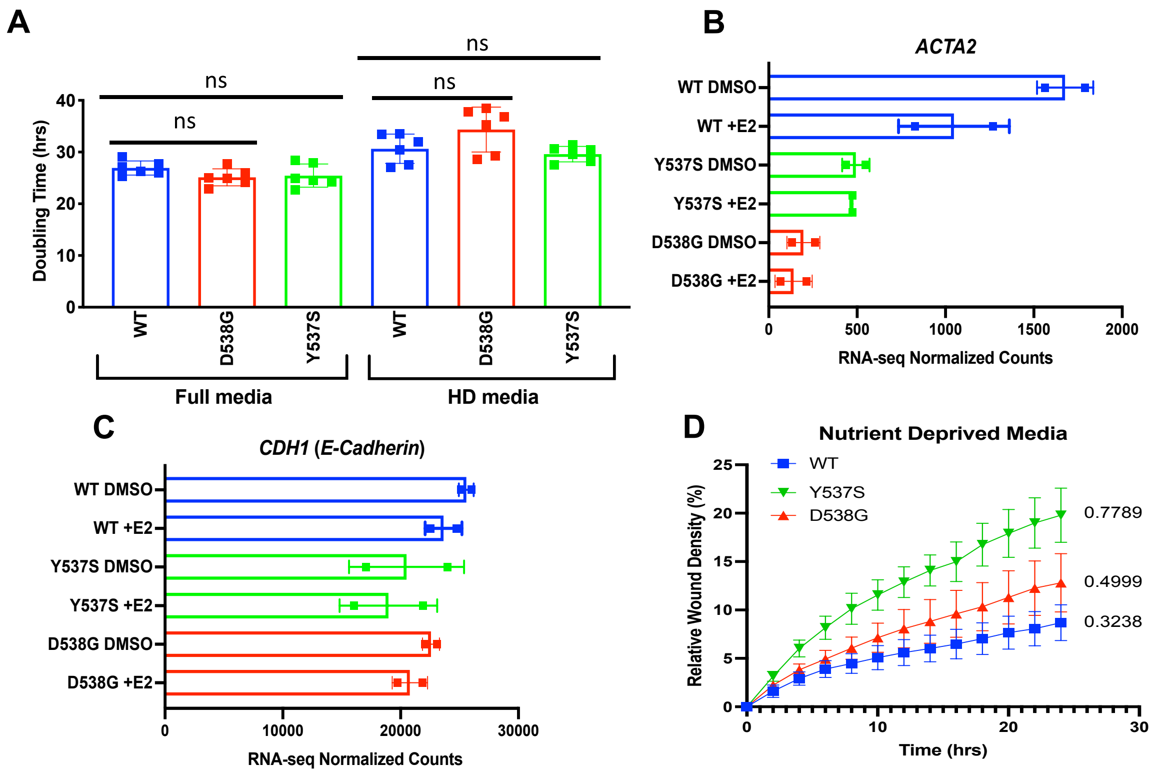


**Supplemental Figure 4. ER mutants do not affect proliferation, but down-regulate proteins associated with movement and impact migration (related to Figure 4).** (A) Proliferation experiments show similar doubling times for ER-WT, ER-D538G, and ER-Y537S cell lines in full serum media and hormone depleted (HD) media. RNA-seq normalized counts of ACTA2 (B) and CDH1 (C). (D) Rates of migration as measured by the relative wound densities of ER-WT, ER-D538G, and ER-Y537S cell lines at 24 hours after performing the scratch assay in nutrient deprived media.

**A**

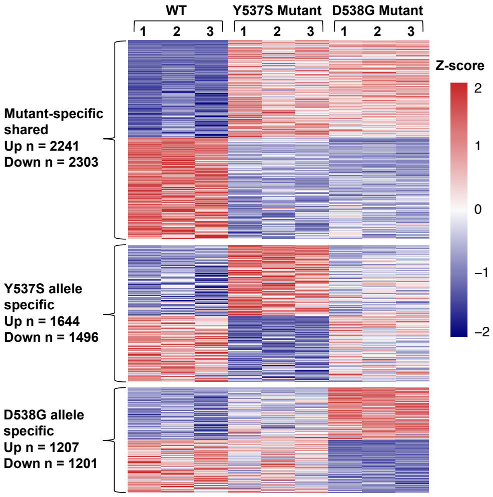


**B**

| Molecular and Cellular Functions [Shared Mutant Genes] | Direction | p-value |
|---|---|---|
| mitotic cell cycle phase transition | ↑ | 2.59E-09 |
| DNA metabolic process | ↑ | 4.58E-09 |
| DNA replication | ↑ | 1.19E-08 |
| G2/M transition of mitotic cell cycle | ↑ | 5.44E-07 |
| G1/S transition of mitotic cell cycle | ↑ | 1.12E-06 |
| double-strand break repair via homologous recombination | ↑ | 1.62E-06 |
| regulation of transcription involved in G1/S transition of mitotic cell cycle | ↑ | 3.13E-06 |
| microtubule bundle formation | ↓ | 2.2103E-05 |

| Molecular and Cellular Functions [D538G Genes] | Direction | p-value |
|---|---|---|
| regulation of transcription, DNA-templated | ↑ | 2.75E-11 |
| regulation of transcription by RNA polymerase II | ↑ | 4.36E-07 |
| double-strand break repair via homologous recombination | ↑ | 4.56E-06 |
| DNA replication | ↑ | 2.29E-05 |
| microtubule cytoskeleton organization involved in mitosis | ↑ | 2.61E-05 |
| DNA repair | ↑ | 5.74E-05 |
| DNA metabolic process | ↑ | 6.63E-05 |
| histone H2A acetylation | ↓ | 1.96E-05 |
| histone H4 acetylation | ↓ | 4.73E-04 |
| regulation of protein phosphorylation | ↓ | 2.81E-03 |

| Molecular and Cellular Functions [Y537S Genes] | Direction | p-value |
|---|---|---|
| translation | ↑ | 4.77E-14 |
| nuclear-transcribed mRNA catabolic process, nonsense-mediated decay | ↑ | 2.48E-12 |
| cellular protein metabolic process | ↑ | 4.89E-10 |
| aerobic electron transport chain | ↑ | 6.60E-10 |
| mitochondrial ATP synthesis coupled electron transport | ↑ | 9.19E-10 |
| de novo' posttranslational protein folding | ↑ | 3.76E-07 |
| rRNA processing | ↑ | 1.49E-06 |
| calcium-dependent cell-cell adhesion via plasma membrane cell adhesion molecules | ↓ | 1.06E-03 |
| cell-cell adhesion via plasma-membrane adhesion molecules | ↓ | 1.94E-03 |
| cell junction assembly | ↓ | 3.66E-03 |

**C**

| MSigDB Hallmark 2020 [Shared Mutant Genes] | Direction | p-value |
|---|---|---|
| E2F Targets | ↑ | 1.83E-18 |
| G2-M Checkpoint | ↑ | 4.46E-12 |
| Estrogen Response Early | ↑ | 2.25E-07 |
| Estrogen Response Late | ↑ | 5.76E-07 |
| mTORC1 Signaling | ↑ | 5.76E-07 |
| Androgen Response | ↑ | 3.13E-06 |
| Epithelial Mesenchymal Transition | ↓ | 7.59E-03 |
| p53 Pathway | ↓ | 7.59E-03 |

| MSigDB Hallmark 2020 [D538G Genes] | Direction | p-value |
|---|---|---|
| E2F Targets | ↑ | 8.77E-13 |
| G2-M Checkpoint | ↑ | 1.54E-11 |
| Mitotic Spindle | ↑ | 4.49E-03 |
| Myc Targets V2 | ↓ | 7.34E-03 |
| Epithelial Mesenchymal Transition | ↓ | 9.10E-03 |

| MSigDB Hallmark 2020 [Y537S Genes] | Direction | p-value |
|---|---|---|
| Oxidative Phosphorylation | ↑ | 2.19E-15 |
| mTORC1 Signaling | ↑ | 3.84E-14 |
| Hypoxia | ↑ | 8.91E-07 |
| Myc Targets V1 | ↑ | 9.01E-05 |
| PI3K/AKT/mTOR Signaling | ↑ | 8.53E-04 |
| Estrogen Response Early | ↑ | 3.84E-03 |

**D**

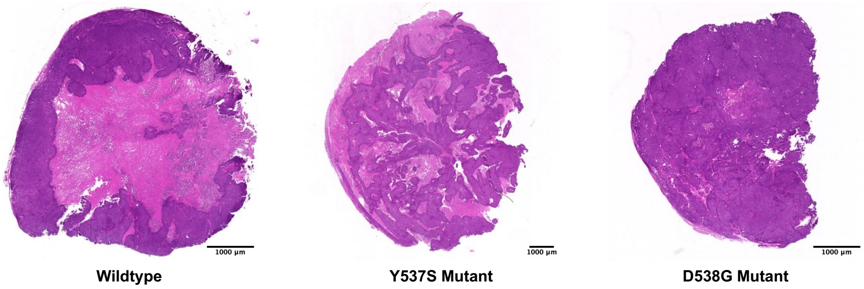


**Supplemental Figure 5. ER-Y537S and ER-D538G mutants drive proliferative gene expression signatures *in vivo* (related to Figure 5).** (A) Heatmap displays relative expression levels of mutant-specific shared, ER-Y537S specific, and ER-D538G specific genes. (B) Enrichr analysis highlights the molecular and cellular functions enriched in genes up- and down-regulated by both ER-D538G and ER-Y537S mutations (left), specific to ER-D538G (middle), and specific to ER-Y537S (right). (C) Enrichr analysis shows MSigDB Hallmark cancer pathways enriched in genes up- and down-regulated by both ER-D538G and ER-Y537S mutations (left), specific to ER-D538G (middle), and specific to ER-Y537S (right). (D) H&E staining is shown for wildtype, ER-Y537S, and ER-D538G xenograft tumors grown in the absence of supplemental estrogen. The wildtype tumor had extensive necrosis in the center of the tumor, the ER-Y537S had mottled necrosis throughout the tumor, and the ER-D538G had relatively little necrosis. Scale bar represents 1000 μm.

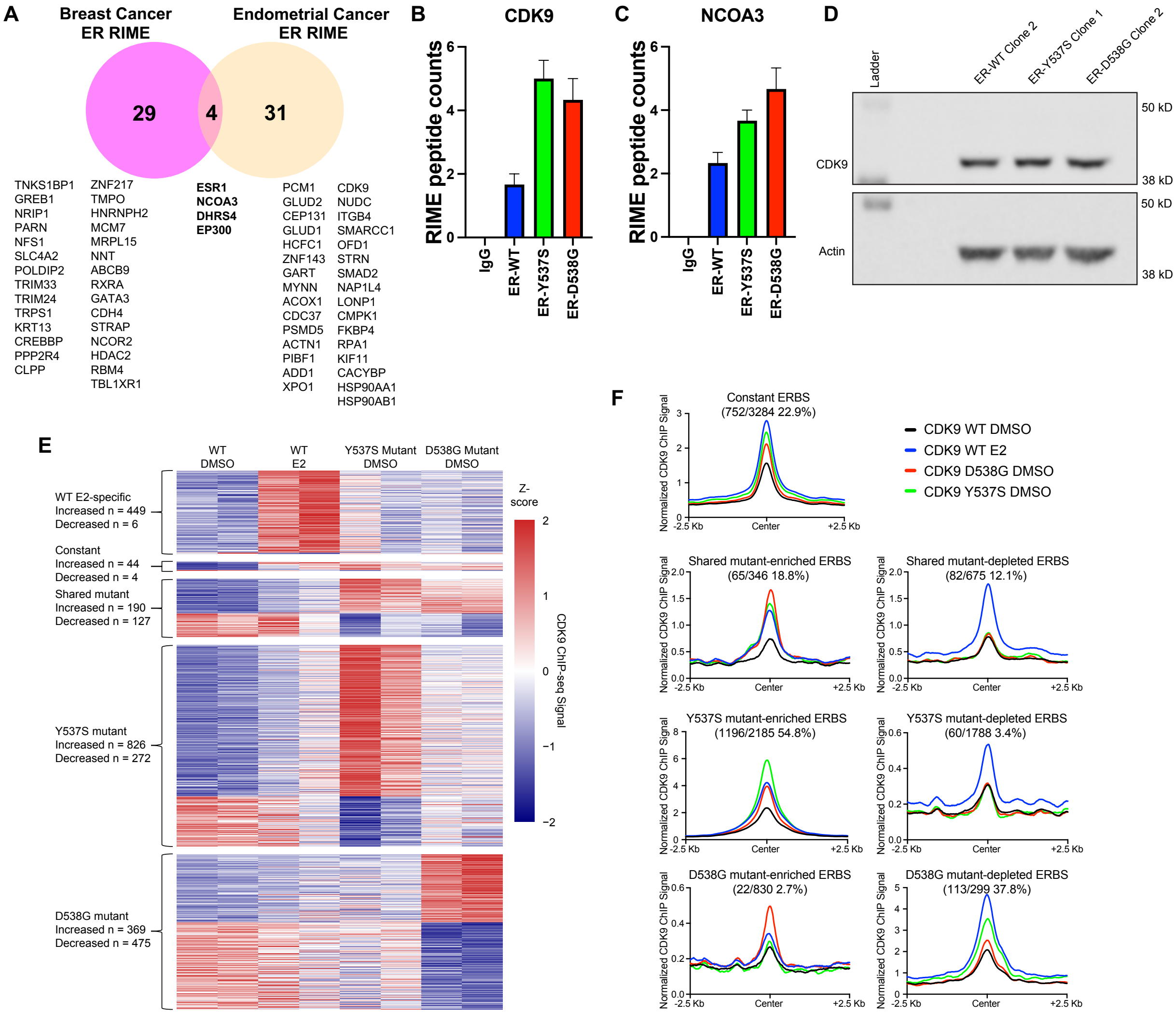


**Supplemental Figure 6. Identification of novel therapeutic targets in ER active and mutant endometrial cancer (related to Figure 6).** (A) Venn diagram shows RIME data highlighting breast and endometrial cancer ER co-associating proteins in MCF-7 (pink) and Ishikawa cell lines (peach). Proteins shared between cancer types are in bold. RIME peptide counts for wildtype ER, ER-Y537S, and ER-D538G are shown for CDK9 (B) and NCOA3 (C). Immunoblotting in Ishikawa ER-WT, ER-Y537S, and ER-D538G clones show protein levels of CDK9 and Actin. (E) Heatmap displays CDK9 ChIP-seq signal at regions enriched or depleted specific to WT with E2 induction, constant in WT with E2 induction and ER LBD mutants with DMSO, shared by both ER LBD mutants, specific to ER-Y537S, and specific to ER-D538G. (F) Aggregate profile plots indicate average signal of CDK9 ChIP-seq at indicated ER binding site (ERBS) class. The number of overlapping CDK9 peaks with ERBS is shown as a fraction of (CDK9 bound sites)/(ERBS) and as a corresponding percentage.

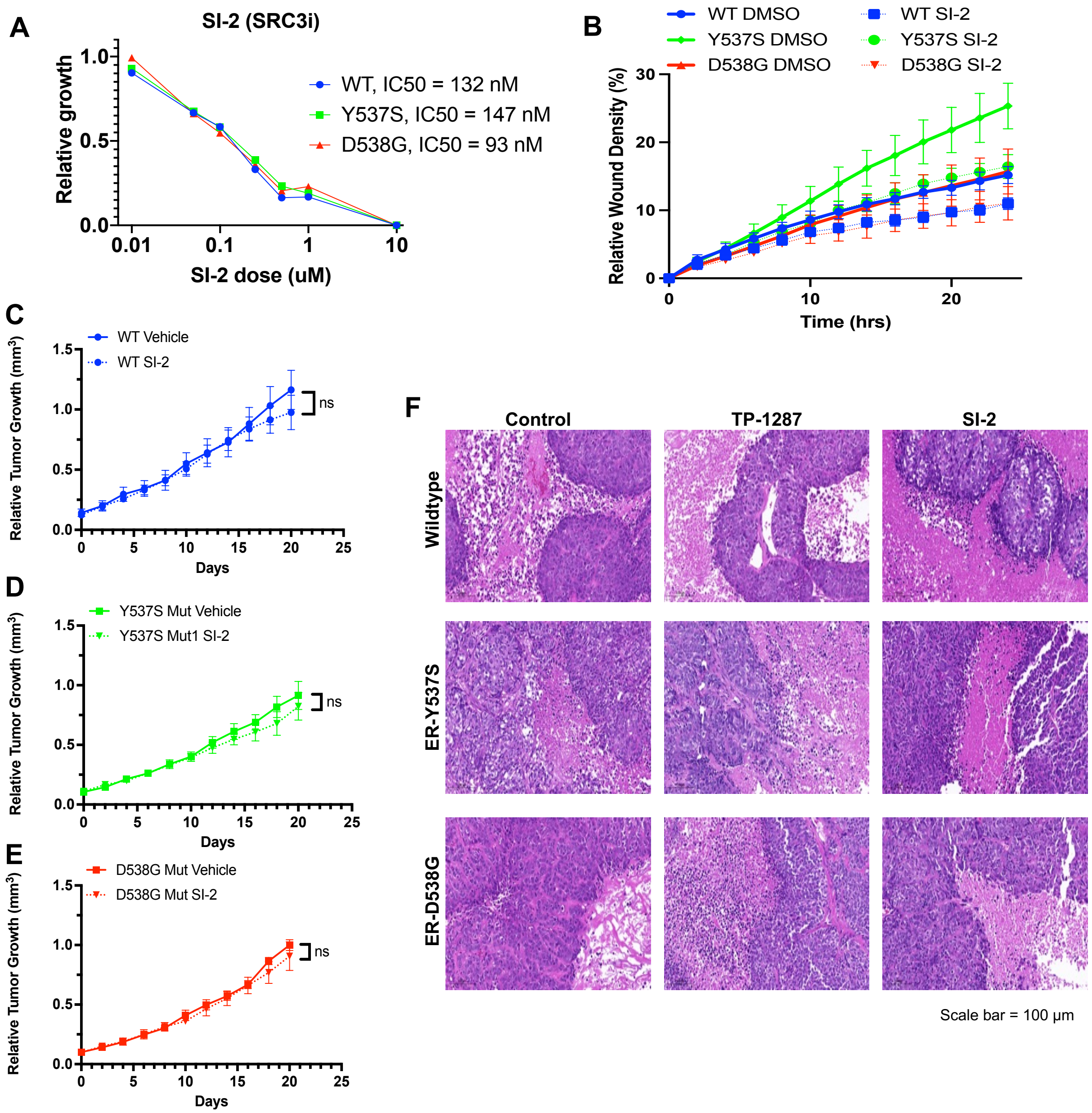


**Supplemental Figure 7. *In vivo* exploration of novel therapeutic targets in ER active and mutant endometrial cancer (related to Figure 6).** (A) Relative growth of wildtype, ER-Y537S, and ER-D538G cell lines after treatment with SI-2 (SRC3 inhibitor) for 72 hours is shown. (B) Reduced rates of migration are observed after SRC3 inhibition, as measured by the relative wound scratch density of wildtype, ER-D538G, and ER-Y537S cell lines in full serum media. Inhibition of SI-2 did not significantly reduce tumor burden in wildtype (C), ER-Y537S (D), and ER-D538G (E) xenograft models. Mice were ovarectomized and received estrogen supplementation. Untreated controls n = 10 mice/group, treated mice n = 8 mice/group. (F) H&E staining is shown for control, TP-1287 (CDK9) and SI-2 (SRC3) treated wildtype, ER-Y537S and ER-D538G tumors. All tumors retained endometrioid histology but differed in degree of necrosis. Scale bar represents 100 uM.

### Supplemental Table 4

Table S4, Related to Methods – Mutant ESR1 generation: gBlock used for template amplification.

| Name | Sequence |
|---|---|
| gBlock (IDT) with Y537S mutation in red | CTTGAACTGCTTTACTCATTTAAAATACCCACTCCTGCTTG GCTGAATATCTCATGTTGTCTTTTTAGAAGCTTTGGCGATC CTATTTGAATGCATTTAGGTCCTATTGGAGGGGAATAGGA TCTCATTTGAGGCCACGGAGGTCCATGGAAGTCACCTGCATA GCAAATACCCTGAAAGTGGCTGCAGGGAGAGTGTGAGGGTG GGACCGCCCTGGTAGGAGGTGGAAAATGAAAAACACACGG CCATGAGTTCCAGATTAGGGCTTCTGAAAGCCCTCAGCTTT CCCAGCTCCCATCCTAAAGTGGGTCTTTAAACAGGAAGAA AGAAAGATTGCTAAGTGTCTTTGGAGTTCCTCTTCCTTCCC CTTCTAGGGATTTCAGCACTCCTGGGGCTCGGGTTGGCTC TAAAGTAGTCCTTTCTGTGTCTTCCCACCTACAGTAACAAA GGCATGGAGCATCTGTACAGCATGAAGTGCAAGAACGTGG TGCCCCTCT**C**TGACCTGCTGCTGGAGATGCTGGACGCCCA CCGCCTACATGCGCCCACTAGCCGTGGAGGGGCATCCGTG GAGGAGACGGACCAAAGCCACTTGGCCACTGCGGGCTCTA CTTCATCGCATTCCTTGCAAAAGTATTACATCACGGGGGAGG CAGAGGGTTTCCCTGCCACGGTCTGAGAGCTCCCTGGCTCC CACACGGTTCAGATAATCC |

### Supplemental Table 5

Table S5, Related to RIME Experiments – Enrichment values for ER associating proteins in endometrial and breast cancer cell lines. Shared proteins are in **bold**. Mutant ER key: S = shared with wildtype ER, Y = ER-Y537S and wildtype ER, D = ER-D538G and wildtype ER, Yup = higher in ER-Y537S than wildtype ER, Dup = higher in ER-D538G than wildtype ER.

| Protein | Endometrial Log2FC | Endometrial Adj P-value | Breast Log2FC | Breast Adj P-value |
|---|---|---|---|---|
| **ESR1** [S] | **5.19** | **2.12E-17** | **4.04** | **1.05E-07** |
| **NCOA3** [S] | **2.12** | **0.02529709** | **3.85** | **0.00254323** |
| **EP300** [S, Yup] | **2.36** | **0.01497222** | **2.66** | **0.03845727** |
| **DHRS4** [S] | **2.34** | **0.00816874** | **3.05** | **0.00147784** |
| XPO1 | 2.13 | 0.03068549 | 1.55 | 0.14558558 |
| HSP90AA1 [D] | 1.6 | 0.00724172 | NA | NA |
| HSP90AB1 [D] | 1.5 | 0.01698495 | NA | NA |
| CDC37 [S] | 2.6 | 0.00033287 | 1.75 | 0.09337269 |
| FKBP4 [S] | 1.87 | 0.00609107 | NA | NA |
| CDK9 [S, Yup, Dup] | 2.1 | 0.04214162 | 1.67 0.06274636 | |
| SMAD2 [S, Yup, Dup] | 2 | 0.04677184 | NA | NA |
| SMARCC1 [S, Yup] | 2 | 0.04677184 | 1.3 | 0.15361547 |
| PCM1 [S] | 6.02 | 9.71E-25 | NA | NA |
| GLUD2 [S] | 5.12 | 1.05E-16 | NA | 0.27167696 |
| CEP131 [S] | 4.97 | 5.84E-16 | NA | NA |
| GLUD1 [Y] | 4.73 | 1.26E-18 | NA | NA |
| HCFC1 [S] | 2.94 | 0.00033287 | NA | NA |
| ZNF143 [S] | 2.94 | 0.00033287 | 0.36 | 0.43313042 |
| GART [S] | 2.85 | 0.00071055 | NA | NA |
| MYNN [S, Yup] | 2.79 | 0.00062781 | NA | NA |
| ACOX1 [Y] | 2.71 | 0.00067504 | NA | NA |
| PSMD5 [S] | 2.45 | 0.00516715 | 1.45 | 0.13829008 |
| ACTN1 | 2.34 | 0.00816874 | 2.06 | 0.09157232 |
| PIBF1 [Y] | 2.23 | 0.01497222 | NA | NA |
| ADD1 [S] | 2.23 | 0.01497222 | NA | NA |
| NUDC [Y] | 2.09 | 0.02298255 | NA | NA |
| ITGB4 [S] | 2 | 0.02081852 | NA | NA |
| ODF1 [S] | 2 | 0.04677184 | NA | NA |
| STRN [S] | 2 | 0.04677184 | NA | NA |
| NAP1L4 [S] | 1.97 | 0.00516715 | 0.6 | 0.36089677 |
| LONP1 | 1.92 | 0.01698495 | 0.8 NA | |
| CMPK1 [S] | 1.91 | 0.03401786 | NA | NA |

| | | | | |
|---|---|---|---|---|
| RPA1 [S] | 1.85 | 0.04677184 | 1.09 | 0.17919395 |
| KIF11 [S] | 1.79 | 0.01109121 | NA | NA |
| CACYBP [S] | 1.65 | 0.02081852 | 1.08 | 0.31056266 |
| CREBBP | 2.01 | 0.06319703 | 3.04 | 0.00258288 |
| GREB1 | NA | NA | 4.5 | 1.55E-08 |
| NCOR2 | NA | NA | 2.34 | 0.02202279 |
| HDAC2 | 0.56 | NA | 2.28 | 0.03084363 |
| TNKS1BP1 | NA | NA | 5.06 | 2.31E-18 |
| GATA3 | NA | NA | 2.5 | 0.01219677 |
| NRIP1 | NA | NA | 4.23 | 0.01900612 |
| PARN | NA | NA | 4.11 | 1.90E-06 |
| NFS1 | NA | NA | 4 | 2.54E-07 |
| SLC4A2 | NA | NA | 3.94 | 0.00014407 |
| POLDIP2 | NA | NA | 3.76 | 2.33E-05 |
| TRIM33 | NA | NA | 3.75 | 0.02808014 |
| TRIM24 | 1.12 | 0.61908834 | 3.48 | 0.00624229 |
| TRPS1 | NA | NA | 3.33 | 0.00014616 |
| KRT13 | -0.47 | NA | 3.25 | 0.0144008 |
| PPP2R4 | NA | NA | 2.97 | 0.01228548 |
| CLPP | -0.87 | 0.79634346 | 2.89 | 0.00491318 |
| ZNF217 | NA | NA | 2.83 | 0.02908564 |
| TMPO | -0.04 | 0.99825043 | 2.79 | 0.01987011 |
| HNRNPH2 | -0.35 | 0.95051335 | 2.78 | 0.01308061 |
| MCM7 | 0.49 | 0.95059486 | 2.75 | 0.00821532 |
| MRPL15 | NA | NA | 2.63 | 0.02856067 |
| NNT | NA | NA | 2.59 | 0.03112125 |
| ABCB9 | NA | NA | 2.53 | 0.02357325 |
| RXRA | NA | NA | 2.51 | 0.04117141 |
| CHD4 | 1.46 | 0.10046436 | 2.44 | 0.0374998 |
| STRAP | 0.31 | 0.97931696 | 2.37 | 0.02976313 |
| RBM4 | 0.25 | 0.99150104 | 2.1 | 0.03345458 |
| TBL1XR1 | 0.13 | NA | 2.08 | 0.04693589 |
